# Pathophysiological Significance of Impaired KAT7-Dependent Histone H3K14 Acetylation During Zinc Deficiency

**DOI:** 10.1101/2023.10.18.562865

**Authors:** Takao Fujisawa, Satoshi Takenaka, Lila Maekawa, Toshiyuki Kowada, Toshitaka Matsui, Shin Mizukami, Yugo Kato, Michio Suzuki, Hisashi Noma, Isao Naguro, Hidenori Ichijo

## Abstract

Zinc is an indispensable micronutrient for optimal physiological functions, and zinc deficiency has been implicated in the pathogenesis of various human diseases. One potential mechanism underlying such pathogenic effects is the alteration of gene expression caused by zinc deficiency; however, the details of this process remain largely unexplored. Here, we show that during zinc deficiency, the histone acetyltransferase KAT7 loses its enzymatic activity, leading to the attenuated acetylation of histone H3 at Lys14 (H3K14ac) at enhancer regions. Physiologically, the decrease in H3K14ac leads to the upregulation of the expression of ZIP10, a plasma membrane-localized zinc transporter, thereby facilitating the import of extracellular zinc to maintain cellular zinc homeostasis. Moreover, prolonged zinc deficiency in mice induced by a zinc-deficient diet or high-fat diet, accompanied by decreased H3K14ac levels in the liver, upregulated the expression of genes associated with intracellular lipid droplet formation, leading to the accumulation of lipids within liver tissue. Our findings demonstrate that cells respond to zinc deficiency by converting it into an epigenetic signal that drives physiological or pathophysiological biological processes.

## Introduction

Zinc is an essential micronutrient that plays a pivotal role in maintaining the structural integrity and enzymatic activities of numerous proteins. Computational analysis revealed that as much as 10% of the human genome encodes a zinc-binding proteins^1^, suggesting the importance of maintaining zinc homeostasis in the human body.

Nevertheless, zinc deficiency occurs, arising from various factors, such as inadequate zinc intake, impaired zinc absorption, and aging^2^. The clinical manifestations of zinc deficiency in humans were first reported in the 1960s^3,4^, and since then, numerous studies have shown that zinc deficiency is strongly associated with the development of various diseases. The classical symptoms of zinc deficiency include skin diseases, mental disorders, immune dysfunction, and growth retardation^5^. Recently, emerging evidence has highlighted correlations between reduced zinc levels and diseases such as cancer^6–12^, inflammatory bowel disease^13^, and chronic kidney disease^14^. However, the role of zinc in the pathogenesis of these conditions and the underlying molecular mechanisms remain to be fully elucidated.

To overcome zinc-deficient conditions, cells need to import zinc from extracellular sources. The Zrt-/Irt-like protein (ZIP) family is a group of zinc ion transporters^15^. Among the 14 human ZIP family proteins, 10 ZIPs (ZIP1-6, 8, 10, 12, and 14) are thought to be localized on the plasma membrane and to be involved in the import of extracellular zinc. Upon zinc deficiency, the abundance of ZIPs on the plasma membrane increases to facilitate zinc uptake as a stress response to maintain cellular zinc homeostasis. One related mechanism involves the upregulation of gene expression through transcriptional control. For example, our previous research demonstrated that zinc deficiency activates the transcription factor ATF6 via the endoplasmic reticulum (ER) stress response, thereby promoting the transcription of ZIP14^16^. In addition, it has been reported that ZIP10 expression increases during zinc deficiency through transcriptional control mediated by the transcription factor MTF1^17^. However, the detailed molecular mechanisms remain largely unclear.

Histone posttranslational modifications (PTMs) are among the key regulators that govern transcriptional control^18^. Each histone PTM is balanced via the regulation of a myriad of enzymes referred to as “writers” and “erasers”; writers catalyse the formation of specific types of PTMs, and erasers remove the PTMs. Interestingly, multiple writers and erasers possess zinc-binding domains^19^, suggesting the critical role of zinc in regulating histone PTMs. Nevertheless, little is known about the dynamics of these writers and erasers during zinc deficiency.

In this study, we observed a drastic decrease in histone H3K14ac during zinc deficiency. Mechanistically, this decrease was driven by the loss of activity of the zinc-dependent writer KAT7, which catalyses H3K14ac formation. In cultured cells, a decrease in H3K14ac led to the transcriptional upregulation of *ZIP10*, thereby importing zinc from extracellular sources to maintain cellular zinc homeostasis. At the organismal level in mice, chronic zinc deficiency in the liver induced the upregulation of the expression of genes involved in lipid droplet synthesis pathways and promoted lipid accumulation in liver tissue via a reduction in KAT7 activity and H3K14ac.

Additionally, we showed that a high-fat diet led to a decrease in hepatic zinc and H3K14ac levels, suggesting that these reductions may more universally contribute to hepatic lipid accumulation. Our data demonstrate that cells respond to zinc deficiency stress by converting it into an epigenetic signal to drive cellular responses.

## Results

### Global decrease in histone H3K14ac during zinc deficiency

We previously reported that the activation of the transcription factor ATF6 elicits the upregulation of ZIP14 expression in HepG2 cells during zinc deficiency^16^. To investigate the impact of ATF6 on global expression changes in genes other than ZIP14, we examined the presence of ATF6-binding motifs on the promoters of genes whose expression was upregulated under zinc-deficient conditions. The Gene Expression Omnibus (GEO) repository contains four datasets reporting gene expression profiles in various types of cells during zinc deficiency, including a dataset from our laboratory^16,20–22^. We found that, across all the datasets, fewer than 2% of the genes presented an ATF6-binding motif^23^ (ER stress-responsive element, ERSE) within their promoter regions (Fig. S1A). These findings indicate that mechanisms other than the ATF6 axis contribute to the alterations in gene expression provoked by zinc deficiency.

In this study, we focused on the role of histone PTMs, which are critical factors responsible for gene expression changes whose association with zinc deficiency remains unexplored. First, we systematically analysed whether zinc deficiency stress affects the major histone PTMs implicated in gene expression in HEK293A cells (promoter marks: H3K4me3, H4K5ac, and H4K12ac; enhancer marks: H3K4me, H3K14ac, and H3K27ac; gene body marks: H3K36me3 and H3K79me3; and heterochromatin marks: H3K9me3 and H3K27me3). We found that treatment with N,N,N’,N’-tetrakis(2-pyridylmethyl)ethylenediamine (TPEN), a cell-permeable zinc chelator, selectively attenuated the signal for H3K14ac; however, no other marks were affected (Fig. 1A). This reduction in H3K14ac was observed across diverse cellular contexts (Fig. S1B, C); this reduction was also evident when cells were cultured in zinc-deficient medium (Fig. 1B), where the reduction was reversed by the addition of zinc and remained unaltered by the addition of other metals (Fig. 1C, D). Furthermore, the H3K14ac signal was restored in a time-dependent manner by the resupplementation of zinc under conditions of zinc deficiency (Fig. 1E). Collectively, these results suggest that zinc reversibly regulates H3K14ac.

**Fig. 1.**
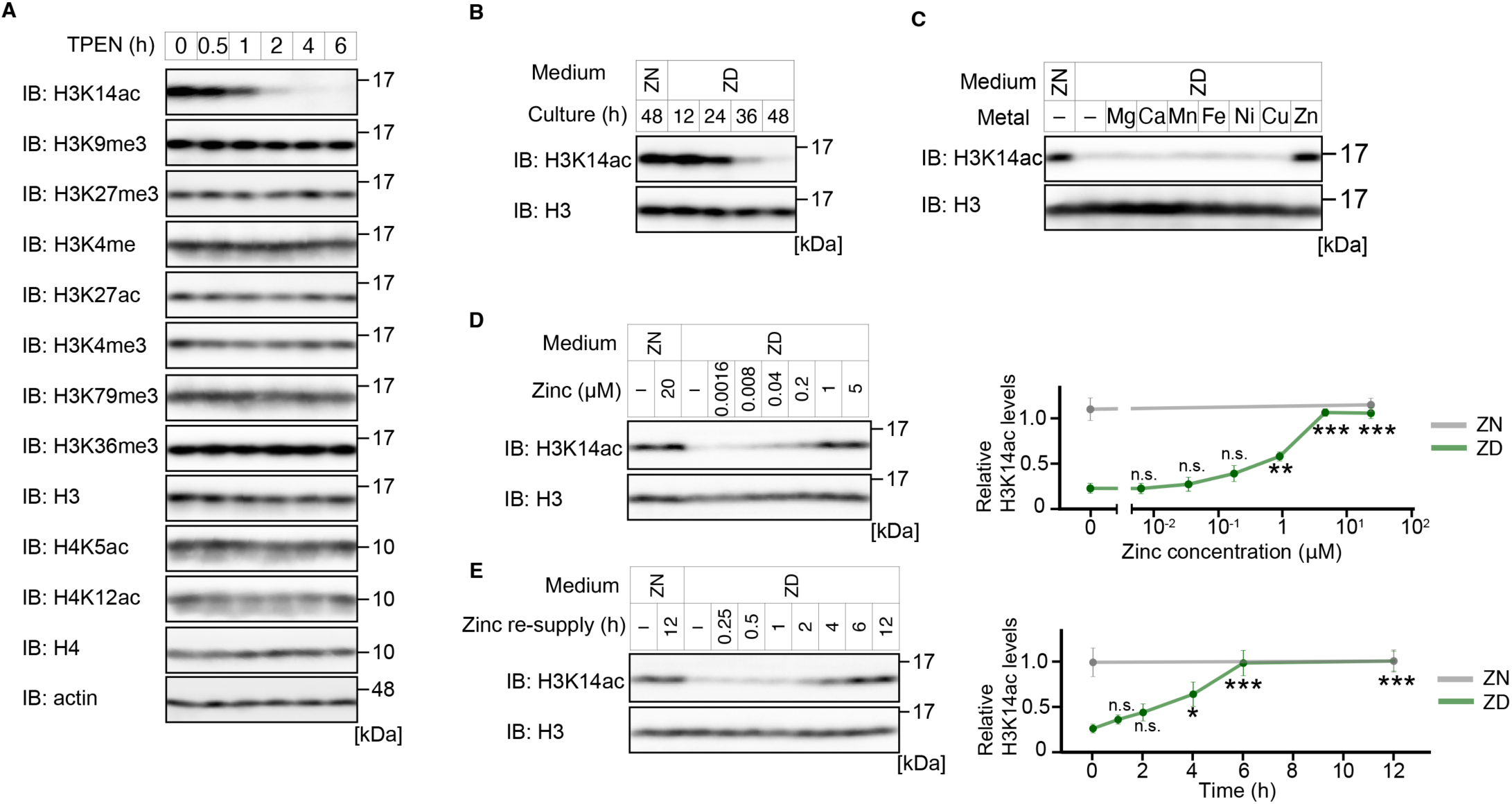
Global decrease in histone H3K14ac during zinc deficiency A) Immunoblot analysis of HEK293A cells treated with TPEN for the indicated times. The data are representative of three independent experiments. B) Immunoblot analysis of HEK293A cells cultured in the indicated medium. The data are representative of three independent experiments. C) Immunoblot analysis of HEK293A cells cultured in the indicated medium supplemented with each metal at 20 µM for 48 hours. The data are representative of three independent experiments. D) Immunoblot analysis of HEK293A cells cultured in zinc-deficient medium supplemented with the indicated concentration of zinc for 48 hours. The data are representative of three independent experiments. The graph shows the mean ± SEM of the H3K14ac level normalized to H3. The H3K14ac level in zinc-containing normal medium was normalized to 1; x-axis, logarithmic scale. The results of statistical analyses comparing the reference group (ZD, no zinc supplementation) to other groups are shown. \*\**P* < 0.01 and \*\*\**P* < 0.001 by one-way ANOVA with Student’s t test followed by Bonferroni post hoc correction. n.s., not significant. E) Immunoblot analysis of HEK293A cells cultured in zinc-deficient medium for 36 hours and additional cultures supplemented with 20 µM zinc for the indicated times. The data are representative of three independent experiments. The graph shows the mean ± SEM of the H3K14ac level normalized to H3. The H3K14ac level in the culture medium was normalized to 1. Data at 0.25 hours and 0.5 hours were omitted from the graph to avoid complexity. The data omitted from the graph were included in the statistical analysis. The results of statistical analyses comparing the reference group (ZD, no zinc resupply) to other groups are shown. \**P* < 0.05 and \*\*\**P* < 0.001 by one-way ANOVA with Student’s t test followed by Bonferroni post hoc correction. n.s., not significant.

### HDACs deacetylate H3K14ac during zinc deficiency

We next investigated the molecular mechanisms underlying the reduction in H3K14ac during zinc deficiency. In general, the histone acetylation status is balanced by histone deacetylases (HDACs) and HATs. We hypothesized that the activation of HDACs or the inactivation of HATs results in a decrease in H3K14ac during zinc deficiency.

First, we investigated the involvement of HDACs. HDACs are classified into two distinct categories: the classic HDAC family (HDAC1-11) and the sirtuin family of NAD-dependent HDACs (SIRT1-7). To identify which HDACs are responsible for the deacetylation of H3K14ac during zinc deficiency, we treated cells with specific inhibitors of each HDAC family. Treatment with nicotinamide (NAM), a paninhibitor of the sirtuin family, failed to inhibit the reduction in H3K14ac (Fig. 2A). In contrast, treatment with trichostatin A (TSA), a paninhibitor of the HDAC family, completely prevented this decrease (Fig. 2B), suggesting that the classic HDAC family deacetylates H3K14ac during zinc deficiency.

**Fig. 2.**
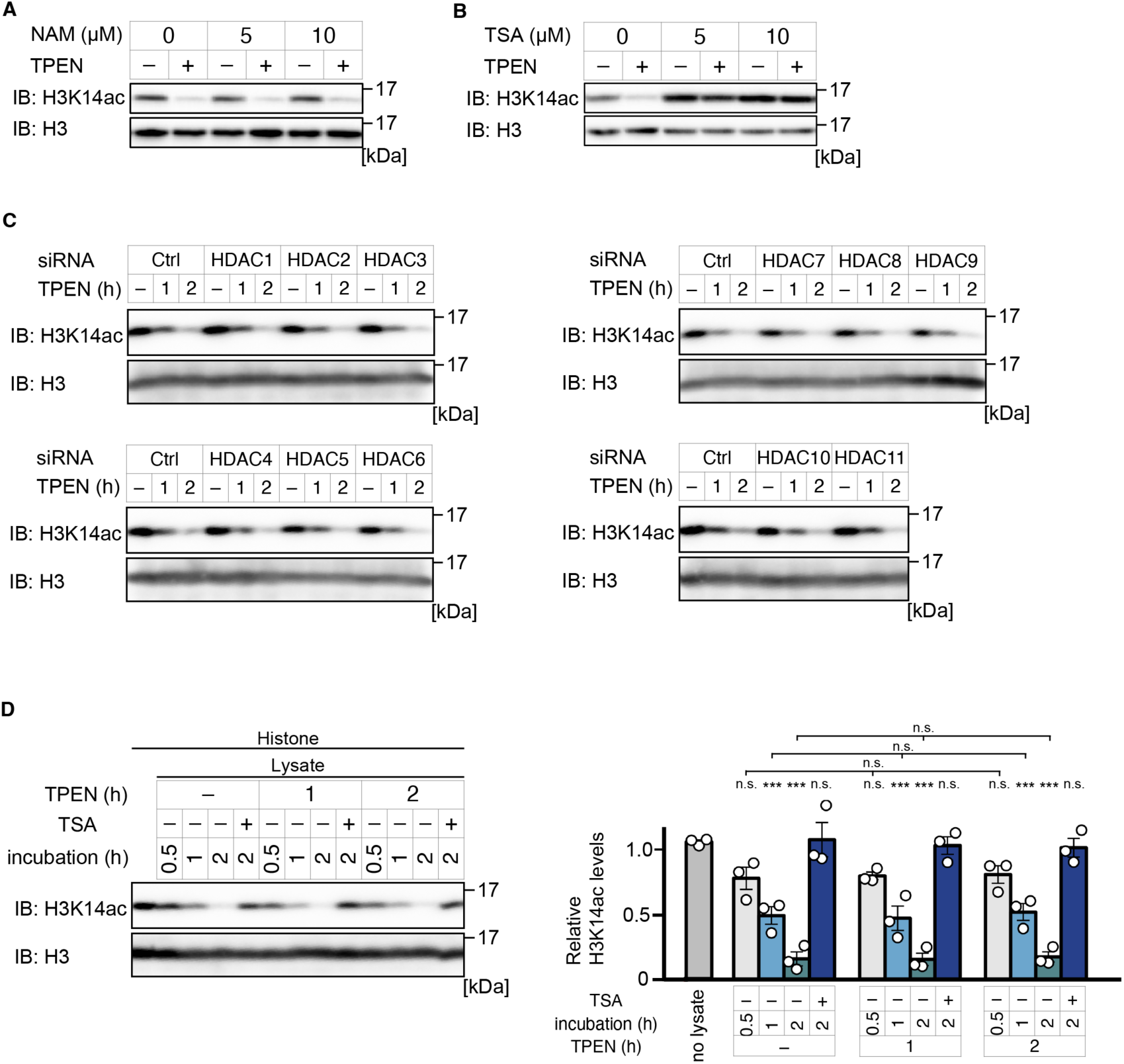
HDACs deacetylate H3K14ac during zinc deficiency A) Immunoblot analysis of HEK293A cells treated with 10 µM TPEN for two hours after pretreatment with NAM for one hour. The data are representative of three independent experiments. B) Immunoblot analysis of HEK293A cells treated with 10 µM TPEN for two hours after pretreatment with TSA for one hour. The data are representative of three independent experiments. C) Immunoblot analysis of HEK293A cells. The cells were treated with the indicated siRNA and cultured for 48 hours. The cells were treated with 10 µM TPEN for the indicated times and subjected to immunoblot analysis. The data are representative of three independent experiments. D) *In vitro* HDAC assay. Cell lysates treated with 10 µM TPEN for the indicated times were prepared. The lysates were incubated with purified histones with or without 2 nM TSA for the indicated times. The data are representative of three independent experiments. The graph shows the mean ± SEM of the H3K14ac level normalized to H3. The H3K14ac level in the absence of cell lysate was normalized to 1. The results of statistical analyses comparing the reference group (no lysate) to other groups are shown on the bars. The results of statistical analyses comparing data from the same incubation times are also shown. \*\*\**P* < 0.001 by one-way ANOVA with Student’s t test followed by Bonferroni post hoc correction. n.s., not significant.

Next, we attempted to identify the specific member of the HDAC family responsible for the deacetylation of H3K14ac during zinc deficiency. However, the knockdown of individual members of the HDAC family did not prevent the TPEN-induced deacetylation of H3K14ac (Fig. 2C). In addition, SAHA, a paninhibitor of the HDAC family, but not other subtype-specific HDAC inhibitors (TMP195: HDAC4/5/7/9; tubastatin: HDAC6; UF010: HDAC1/2/3/6/8/10; MS275: HDAC1/2/3/8) effectively inhibited the TPEN-induced deacetylation of H3K14ac (Fig. S2A, B). These data suggest that several HDAC family members cooperatively contribute to the deacetylation of H3K14ac during zinc deficiency.

To examine the possibility that zinc-deficient stress enhances the deacetylation activity of members of the HDAC family towards H3K14ac, we measured HDAC activity via an *in vitro* HDAC assay using purified histones. The basal HDAC activity for H3K14ac in cell lysates did not differ in the presence or absence of TPEN, suggesting that zinc deficiency in cells enhances the deacetylation of H3K14ac through an independent mechanism that regulates HDAC activity (Fig. 2D). These data suggest that certain members of the classic HDAC family are needed to deacetylate H3K14ac during zinc deficiency but are not direct regulatory targets of zinc deficiency.

### Loss of KAT7 activity during zinc deficiency

Given that HDAC activity is not enhanced under zinc-deficient conditions, we hypothesized that zinc deficiency suppresses the activity of HATs towards H3K14. First, to identify the specific HAT responsible for H3K14ac in HEK293A cells, we performed an siRNA screen using siRNAs targeting 17 human HATs and found that the knockdown of K (lysine) acetyl transferase 7 (KAT7) led to a significant decrease in H3K14ac (Fig. 3A). This decrease was also observed when the cells were treated with WM-3835, a specific inhibitor of KAT7^24^ (Fig. S3A), indicating the crucial role of KAT7 in the acetylation of H3K14. Thus, we focused on the regulatory mechanisms of the KAT7 HAT activity during zinc deficiency.

**Fig. 3.**
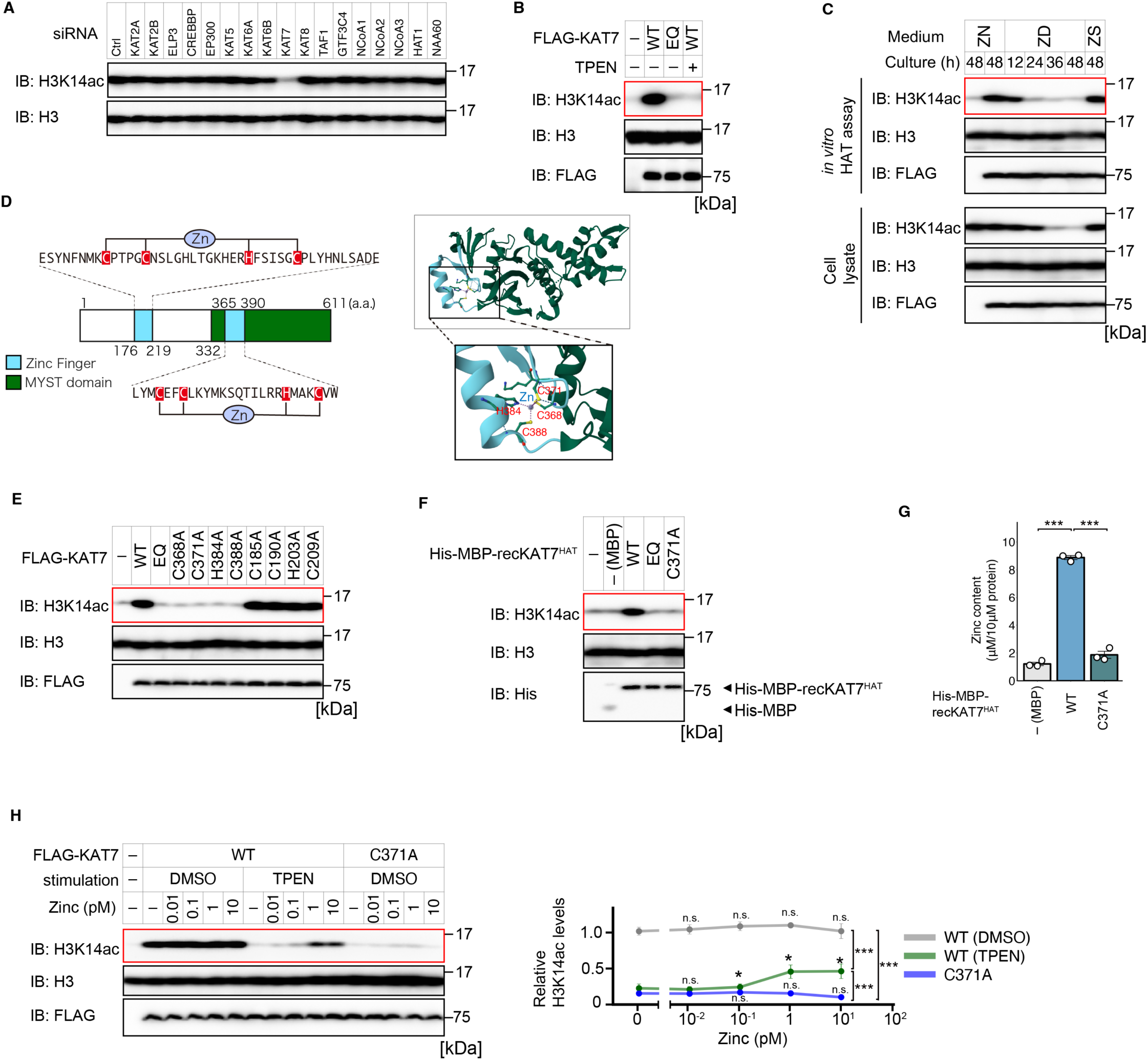
Zinc coordination in the KAT7 MYST domain regulates HAT activity towards H3K14 A) Immunoblot analysis of HEK293A cells treated with the indicated siRNAs for 48 hours. The data are representative of three independent experiments. B) *In vitro* HAT assay. FLAG-KAT7 immunoprecipitated from HEK293A cells transfected with the indicated plasmids and treated with or without 10 µM TPEN for two hours was subjected to an *in vitro* HAT assay. The data are representative of three independent experiments. C) Upper panels, *in vitro* HAT assay. FLAG-KAT7 WT immunoprecipitated from HEK293A cells cultured in the indicated medium for 48 hours was subjected to an *in vitro* HAT assay. Lower panels, immunoblot analysis of HEK293A cells transfected with FLAG-KAT7 WT and incubated with the indicated medium. ZN, zinc-normal medium; ZD, zinc-deficient medium; ZS, zinc-deficient medium supplemented with 20 µM ZnCl2. The data are representative of three independent experiments. D) Left, schematic representation of the KAT7 domain structure and amino acid sequences in zinc finger domains. Right, 3D structures of the KAT7 MYST domain (Protein Data Bank, 6MAJ). Blue, zinc finger domain. The amino acids coordinating zinc are highlighted. E) *In vitro* HAT assay. FLAG-KAT7 WT or mutants immunoprecipitated from HEK293A cells were subjected to an *in vitro* HAT assay. The data are representative of three independent experiments. F) *In vitro* HAT assay. RecKAT7^HAT^ with or without mutation was subjected to a HAT assay. The data are representative of three independent experiments. G) ZnAF-2 assay. The zinc content in recKAT7^HAT^ was measured. The graph shows the mean ± SEM (n=3). \*\*\**P* < 0.001 by one-way ANOVA with Student’s t test followed by Bonferroni post hoc correction. n.s., not significant. H) *In vitro* HAT assay. FLAG-KAT7 WT or C371A immunoprecipitated from HEK293A cells treated with or without 10 µM TPEN for two hours was incubated with the indicated concentration of ZnCl2 for 30 minutes. The samples were subjected to an *in vitro* HAT assay. The data are representative of three independent experiments. The results of statistical analyses comparing the reference group (with no additional zinc) to other groups are shown on the plots. The results of statistical analyses comparing data collected following 10 pM zinc supplementation are shown. \**P* < 0.05 and \*\*\**P* < 0.001 by one-way ANOVA with Student’s t test followed by Bonferroni post hoc correction. n.s., not significant.

Previous reports have shown that KAT7 undergoes degradation in response to DNA damage and lipopolysaccharide stimulation via the ubiquitin‒proteasome pathway^25,26^. However, our investigation indicated that the effect of TPEN on the protein level of KAT7 was limited (Fig. S3B). While treatment with the proteasome inhibitor MG132, but not the autophagy inhibitor bafilomycin A1, inhibited the TPEN-induced slight decrease in the protein amount of KAT7, both inhibitors had little effect on the decrease in H3K14ac. These results suggest that the degradation of KAT7 is not involved primarily in the reduction in H3K14ac during zinc deficiency.

Several reports have demonstrated that KAT7 localizes to the nucleus to acetylate histones^27,28^. We investigated the possibility that the subcellular localization of KAT7 changes during zinc deficiency. However, immunofluorescence analysis of endogenous KAT7 revealed that KAT7 localized to the nucleus after TPEN treatment (Fig. S3C), suggesting that a change in subcellular localization is not involved in the regulation of KAT7 during zinc deficiency.

Interactome analyses demonstrated that KAT7 forms a multisubunit complex that includes BRPF1/2/3, JADE1/2/3, ING4/5, and MEAF6^29–35^. Given that these interactors have been reported to directly bind to KAT7 to increase its HAT activity, we hypothesized that zinc deficiency leads to the dissociation of KAT7 from the complex. The systematic knockdown of each interactor revealed that BRPF2 and MEAF6 play critical roles in the acetylation of H3K14 in HEK293A cells (Fig. S3D). However, treatment with TPEN did not dissociate KAT7 from each component (Fig. S3E). These findings suggest that the disruption of the KAT7 complex is not a cause of the reduction in H3K14ac during zinc deficiency.

Since working hypotheses based on previous reports were not applicable in this case, we next investigated whether zinc deficiency directly affects KAT7 activity by employing an *in vitro* HAT assay. The treatment of cells with TPEN strongly inhibited the HAT activity of immunoprecipitated KAT7 to a similar level as that of the HAT-dead EQ mutant (Fig. 3B). In addition, the KAT7 HAT activity in cells cultured in low-zinc-medium was attenuated in a time-dependent manner, which was followed by the deacetylation of H3K14ac (Fig. 3C). These results suggest that KAT7 loses its enzymatic activity under zinc deficiency.

### Zinc coordination in the KAT7 MYST domain regulates HAT activity towards H3K14

KAT7 possesses two zinc finger domains: one is located within the MYST-type HAT domain (MYST domain), and the other is in the N-terminal region (Fig. 3D). We speculated that the loss of zinc coordination during zinc deficiency affects KAT7 HAT activity. To test this possibility, we examined whether the zinc finger domains in KAT7 are needed for KAT7 HAT activity. Depleting KAT7 using the CRISPR‒Cas9 system resulted in decreased H3K14ac levels in HEK293A cells (Fig. S3F). Adding back KAT7 wild-type (WT) or KAT7 mutants with mutations in the N-terminal zinc finger (C185A, C190A, H203A, and C209A) into the KAT7 knockout HEK293A cell line restored the H3K14ac level (Fig. S3F). In contrast, the reintroduction of KAT7 mutants with mutations in the zinc finger within the MYST domain (C368A, C371A, H384A, and C388A) failed to do so. In support of that finding, immunoprecipitated KAT7 mutants harbouring a zinc finger mutation in the MYST domain showed no HAT activity for H3K14 *in vitro*, whereas KAT7 mutants with a zinc finger mutation in the N-terminal region presented HAT activity similar to that of WT KAT7 (Fig. 3E). These results suggest that the zinc finger within the MYST domain and zinc-binding amino acid residues are needed for KAT7 HAT activity.

Next, we analysed the properties of recombinant proteins purified from *E. coli*. We attempted to obtain full-length KAT7 but were unsuccessful. Thus, we purified the recombinant KAT7 HAT domain (recKAT7^HAT^), which includes the MYST domain. Consistent with the results above, recKAT7^HAT^ WT, but not the recKAT7^HAT^ EQ mutant or recKAT7^HAT^ C371A mutant, exhibited HAT activity towards H3K14 (Fig. 3F). Notably, a ZnAF-2 assay, a method to quantify the zinc content coordinated within recombinant proteins, demonstrated that recKAT7^HAT^ C371A coordinates with substantially less zinc than recKAT7^HAT^ WT does (Fig. 3G), suggesting the critical role of proper zinc coordination in the MYST domain for KAT7 HAT activity.

To further assess the role of zinc in KAT7, we manipulated the zinc metalation status of KAT7 *in vitro* and examined the effect on HAT activity. First, we attempted to demetallate zinc from KAT7 *in vitro*. However, even when recKAT7^HAT^ WT was treated with TPEN *in vitro*, recKAT7^HAT^ WT still coordinated with zinc and exhibited HAT activity (Fig. S3G, H). Consistent with these findings, *in vitro* TPEN treatment of KAT7 immunoprecipitated from HEK293A cells did not affect HAT activity (Fig. S3I). These results suggest that KAT7 coordinates zinc much more strongly than TPEN does and that the elimination of zinc from KAT7 *in vitro* was unsuccessful. Next, we examined the effect of *in vitro* zinc repletion. As expected, *in vitro* zinc addition to KAT7 WT immunoprecipitated from HEK293A cells treated with TPEN, but not KAT7 C371A, restored the HAT activity in a dose-dependent manner (Fig. 3H). Collectively, these results suggest that KAT7 loses zinc from the MYST-type zinc finger domain during zinc deficiency, which leads to decreased HAT activity towards H3K14.

### Loss of H3K14ac on the enhancer induces *ZIP10* expression to maintain cellular zinc homeostasis

Next, we investigated the physiological role of transcriptional changes through the drastic reduction in H3K14ac during zinc deficiency. We and others have previously shown that H3K14ac regulates gene expression at enhancers^36–38^. Moreover, ChIP-seq analysis revealed a significant decrease in H3K14ac on enhancer regions in HeLa cells under zinc-deficient conditions and upon KAT7 knockdown (Fig. 4A). Thus, we speculated that genes responsible for maintaining cellular zinc homeostasis are regulated by H3K14ac. The Molecular Signatures Database (MSigDB) lists genes involved in zinc homeostasis (GO:0006829, zinc ion transport)^39,40^. We systematically integrated our gene expression profile data with H3K14ac ChIP-seq data under conditions of zinc deficiency and KAT7 knockdown. This analysis revealed that multiple genes were transcriptionally altered, with decreased H3K14ac signals at their enhancer regions under zinc-deficient conditions and upon KAT7 knockdown (Fig. 4B), suggesting that these genes may be regulated by a reduction in H3K14ac. Among these genes, we became interested in plasma membrane-localized ZIPs, which play a key role in importing zinc from extracellular sources. Quantitative PCR (qPCR) analysis results confirmed our previous microarray findings (Fig. 4C). In particular, the notable upregulation of *ZIP10* expression led us to focus on the physiological role of upregulated *ZIP10* expression during zinc deficiency.

**Fig. 4.**
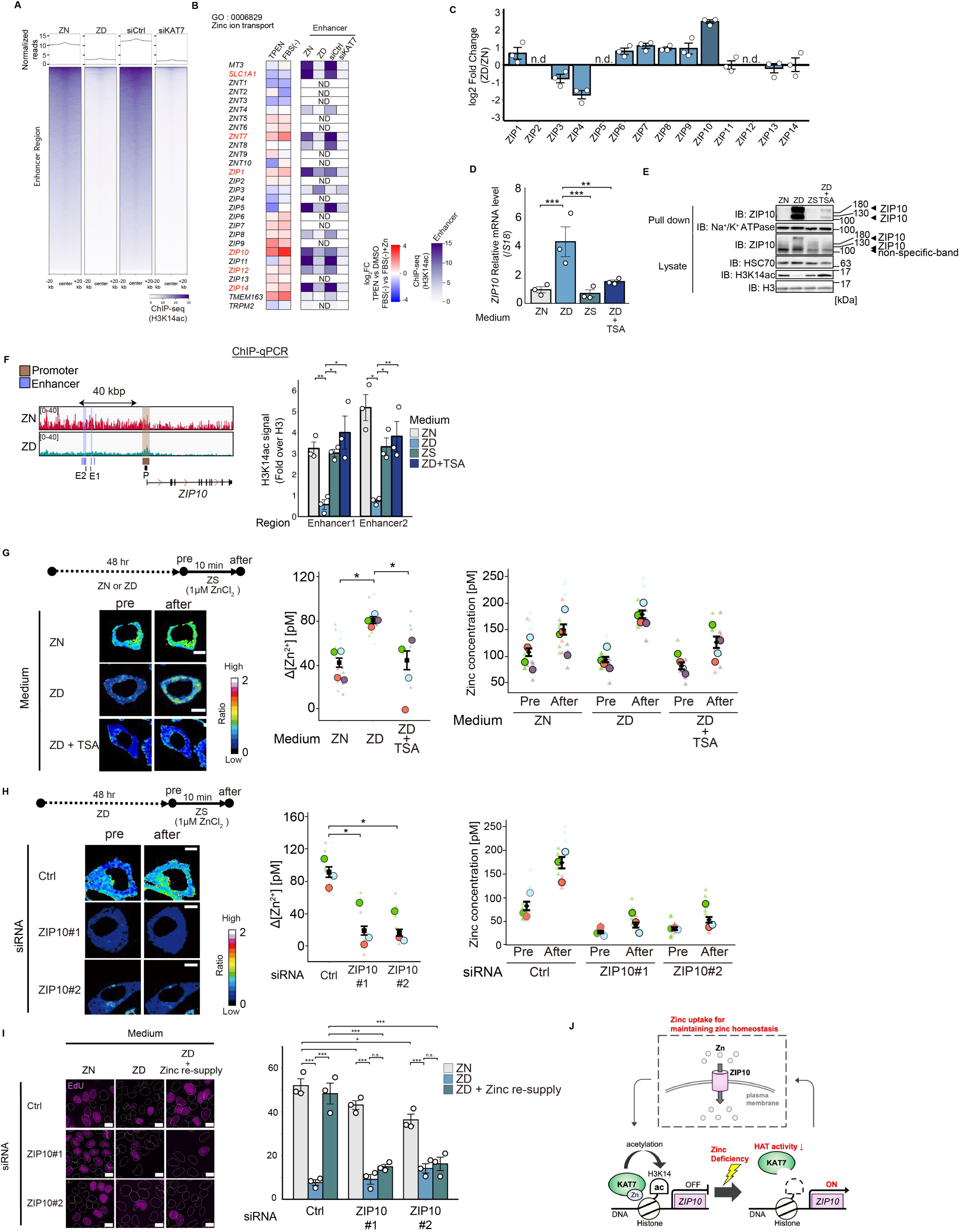
Loss of H3K14ac on the enhancer induces *ZIP10* expression to maintain cellular zinc homeostasis A) Average plot (top) and heatmap (bottom) of H3K14ac ChIP-seq reads around enhancer regions (n=63285). HeLa cells were treated with the indicated siRNA or cultured in the indicated medium for 48 hours. B) Heatmaps showing gene expression changes during zinc deficiency and the H3K14ac signal in enhancer regions. Genes in red are possible candidates regulated by the decrease in H3K14ac during zinc deficiency and upon KAT7 knockdown, as defined by the following criteria: a fold change in expression > 1.5 (upregulation or downregulation, data from GSE49657). ND, not determined (no enhancer information on FANTOM5). C) Quantitative PCR analysis of HeLa cells cultured in the indicated medium for 48 hours. The graph shows the mean ± SEM (n=3). n.d., not detected. D) Quantitative PCR analysis of HeLa cells cultured in the indicated medium supplemented with 10 µM ZnCl2 or 0.5 µM TSA for 36 hours. TSA was added every 12 hours. The data are presented as means ± SEM (n=3). \*\**P* < 0.01 and \*\*\**P* < 0.001 by one-way ANOVA followed by Dunnett’s multiple comparison test. E) Membrane protein biotinylation assay and immunoblot analysis of HeLa cells cultured in the indicated medium supplemented with 10 µM ZnCl2 or 0.5 µM TSA for 48 hours. TSA was added every 12 hours. The data are representative of three independent experiments. Immunoblotting for Na^+^/K^+^ ATPase was performed to determine the amount of membrane proteins. F) Left, H3K14ac ChIP-seq data. The enhancer regions deposited in EnhancerAtlas 2.0 around *ZIP10*, the promoter region of *ZIP10*, and the regions used to design the primers for ChIP‒qPCR are shown. E1, enhancer region 1; E2, enhancer region 2; P, promoter region. Right, ChIP‒qPCR analysis of HeLa cells cultured in the indicated medium supplemented with 10 µM ZnCl2 or 0.5 µM TSA for 36 hours. TSA was added every 12 hours. The graph shows the mean ± SEM (n=3) of the H3K14ac ChIP‒qPCR signal normalized to the H3 ChIP‒qPCR signal. \**P* < 0.05 and \*\**P* < 0.01 by one-way ANOVA with Student’s t test followed by Bonferroni post hoc correction. G) Left, pseudocolour fluorescence ratio images of ZnDA-3H/HTL-TMR. HeLa cells were cultured in the indicated medium for 12 hours. The cells were stimulated with 0.5 µM TSA for 36 hours. TSA was added every 12 hours. The difference in the zinc concentration before and 10 min after the addition of 1 µM zinc is shown. Scale bars, 10 µm. The data are representative of four independent experiments. Middle, labile zinc influx in the cytoplasm measured by ZnDA-3H. A triangle indicates each data point, a circle indicates the mean of each experiment, and a rectangle indicates the mean of all experiments. Right, Labile zinc concentration in the cytoplasm measured by ZnDA-3H. A triangle indicates each data point, a circle indicates the mean of each experiment, and a rectangle indicates the mean of all experiments. The bar represents the mean ± SEM. Data for NZ (n=22), ZD (n=17), and ZD+TSA (n=12) were obtained from 4 independent experiments. pre, before zinc supplementation; after, 10 minutes after zinc supplementation. ZN, zinc-normal medium; ZD, zinc-deficient medium; ZS, zinc-deficient medium supplemented with 10 µM ZnCl2. The bar represents the mean ± SEM. Data for ZN (n=22), ZD (n=17), and ZD+TSA (n=12) were obtained from 4 independent experiments. \**P* < 0.05 by one-way ANOVA followed by Dunnett’s multiple comparison test. H) Left, pseudocolour fluorescence ratio images of ZnDA-3H/HTL-TMR. HeLa cells were treated with the indicated siRNA for 24 hours. The cells were cultured in zinc-deficient medium for an additional 48 hours. The difference in the zinc concentration before and 10 min after the addition of 1 µM zinc is shown. Scale bars, 10 µm. The data are representative of three independent experiments. Middle, labile zinc influx in the cytoplasm measured by ZnDA-3H. A triangle indicates each data point, a circle indicates the mean of each experiment, and a rectangle indicates the mean of all experiments. The bar represents the mean ± SEM. Data for the Ctrl (n=13), ZIP10#1 (n=18), and ZIP10#2 (n=17) groups were obtained from 3 independent experiments. \**P* < 0.05 by one-way ANOVA followed by Dunnett’s multiple comparison test. Right, Labile zinc concentration in the cytoplasm measured by ZnDA-3H. A triangle dot indicates each data point, a circle indicates the mean of each experiment, and a rectangle dot indicates the mean of all experiments. The bar represents the mean ± SEM. Data for the Ctrl (n=13), ZIP10#1 (n=18), and ZIP10#2 (n=17) groups were obtained from 3 independent experiments. pre, before zinc supplementation; after, 10 minutes after zinc supplementation. I) EdU assay. HeLa cells treated with the indicated siRNA were cultured in the indicated medium for 48 hours. After incubation with or without 10 µM ZnCl2 for 9 hours, an EdU assay was conducted. The white lines indicate the boundaries of the nucleus. The data are presented as means ± SEM (n=3). \**P* < 0.05 and \*\*\**P* < 0.001 by one-way ANOVA with Student’s t test followed by Bonferroni post hoc correction. n.s., not significant. J) Schematic of the model reported in this study. ZN, zinc-normal medium; ZD, zinc-deficient medium; ZS, zinc-deficient medium supplemented with 10 µM ZnCl2.

In HeLa cells, zinc supplementation suppressed the decrease in H3K14ac during zinc deficiency, which was strongly correlated with KAT7 HAT activity (Fig. S4A, B). Moreover, treatment with TSA also suppressed the decrease in H3K14ac but did not restore KAT7 HAT activity. Under these experimental conditions, we examined whether the zinc-dependent regulation of H3K14ac regulates *ZIP10* expression. Zinc supplementation or TSA treatment counteracted the upregulation of *ZIP10* expression during zinc deficiency (Fig. 4D). Using a membrane protein biotinylation assay, we confirmed that zinc deficiency led to the expression of *ZIP10* at the plasma membrane, an effect that was suppressed by treatment with zinc or TSA (Fig. 4E). In addition, ChIP-seq and ChIP‒qPCR analyses revealed that the H3K14ac signal in the enhancer region of *ZIP10*, which was predicted by EnhancerAtlas 2.0^41^, significantly decreased during zinc deficiency, whereas the signal in the promoter region did not decrease (Fig. 4F, Fig. S4C). This decrease in the enhancer regions was completely suppressed by treatment with zinc or TSA. Moreover, this decrease in the enhancer region of H3K14ac and increase in *ZIP10* mRNA were dependent on KAT7, as confirmed after KAT7 knockdown (Fig. S4D-F). These results suggest that *ZIP10* expression is regulated by KAT7-catalysed H3K14ac at enhancer regions.

To investigate whether the upregulation of *ZIP10* expression contributes to maintaining zinc homeostasis, we assessed cellular zinc levels indirectly by measuring *MT2A* mRNA, the expression of which is well known to change depending on the concentration of cellular zinc^42,43^. Consistent with previous reports, zinc supplementation drastically upregulated *MT2A* expression in cells cultured in low-zinc medium, an effect that was suppressed by the knockdown of *ZIP10* (Fig. S4G).

Notably, the knockdown of *ZIP10* but not other *ZIPs* significantly attenuated the upregulation of *MT2A* expression (Fig. S4H), suggesting the critical role of ZIP10 in the import of zinc during zinc deficiency. To directly assess zinc uptake through ZIP10, we employed ZnDA-3H, a cell-permeable fluorescent Zn^2+^ probe that can be localized to specific subcellular compartments, via HaloTag technology^44^. ZnDA-3H was localized in the cytosol when Halo-NES was expressed, and zinc influx was measured under these experimental conditions. The import of zinc (Δ[Zn^2+^] = 80.8 ± 3.0 pM) was significantly greater in the cells cultured in the low-zinc medium than in the cells cultured in the normal medium (42.4 ± 4.0 pM) (Fig. 4G). In contrast, TSA treatment suppressed zinc uptake (44.5 ± 8.4 pM), suggesting a critical role for H3K14ac in zinc influx. Notably, zinc uptake was significantly lower after ZIP10 knockdown (siRNA#1, 18.7 ± 5.3 pM; siRNA#2, 16.2 ± 4.3 pM) than after control knockdown (91.9 ± 6.2 pM) (Fig. 4H). Collectively, these data suggest that the zinc influx induced by the addition of zinc to zinc-deficient cells is driven primarily by the H3K14 deacetylation-dependent increase in ZIP10 expression.

Next, we examined the biological role of upregulated ZIP10 expression during zinc deficiency. A previous study showed that zinc deficiency induces cell cycle arrest and that subsequent zinc repletion facilitates cell cycle re-entry^45^. By employing an EdU assay, we confirmed that zinc deficiency indeed arrests the cell cycle and that subsequent zinc supplementation induces cell cycle re-entry (Fig. 4I, siCtrl). Notably, ZIP10 knockdown inhibited the process of cell cycle re-entry (Fig. 4I, siZIP10#1 and siZIP10#2). These findings reinforce the importance of upregulated ZIP10 expression during zinc deficiency in cellular biological processes (Fig. 4J).

### Lipid accumulation in the liver during zinc deficiency

Next, we investigated the biological significance of the zinc deficiency-induced reduction in H3K14ac *in vivo*. Given the central role of the liver in zinc metabolism, we examined the expression of ZIP10 in the livers of mice fed a zinc-deficient diet. Consistent with the findings in HeLa cells, *ZIP10* expression was significantly upregulated (Fig. S5A), suggesting that zinc uptake through the increased expression of ZIP10 may also contribute to maintaining zinc homeostasis *in vivo*. Moreover, if zinc deficiency persists due to a prolonged unbalanced diet, the limited availability of zinc for cellular uptake may disrupt cellular zinc homeostasis, potentially leading to abnormalities in biological functions. Thus, we investigated the possibility that prolonged zinc deficiency contributes to disease development by inducing sustained changes in gene expression associated with reduced H3K14ac levels *in vivo*.

We utilized the gene expression database Expression Atlas^46^ to investigate diseases potentially linked to a reduction in zinc levels. Focusing on the *metallothionein 2* (*Mt2*) gene, whose gene expression changes depending on the cellular zinc level, we examined the gene expression profile datasets of the mice with reduced *Mt2* expression. There were 174 datasets in which *Mt2* expression was significantly decreased. Among them, we focused on the "control vs. high-fat diet" data for the liver, which reported the most significant decrease in *Mt2* expression (Fig. 5A). On the basis of these publicly available data, a high-fat diet in mice may lead to a significant reduction in hepatic zinc levels. This finding was confirmed by inductively coupled plasma‒mass spectrometry (ICP‒MS) analysis, which revealed a marked reduction in hepatic zinc concentrations in mice fed a high-fat diet (Fig. 5B). Since a high-fat diet induces hepatic lipid accumulation and promotes the development of fatty liver, we investigated the possibility that the reduction in liver zinc levels is involved in hepatic steatosis.

**Fig. 5.**
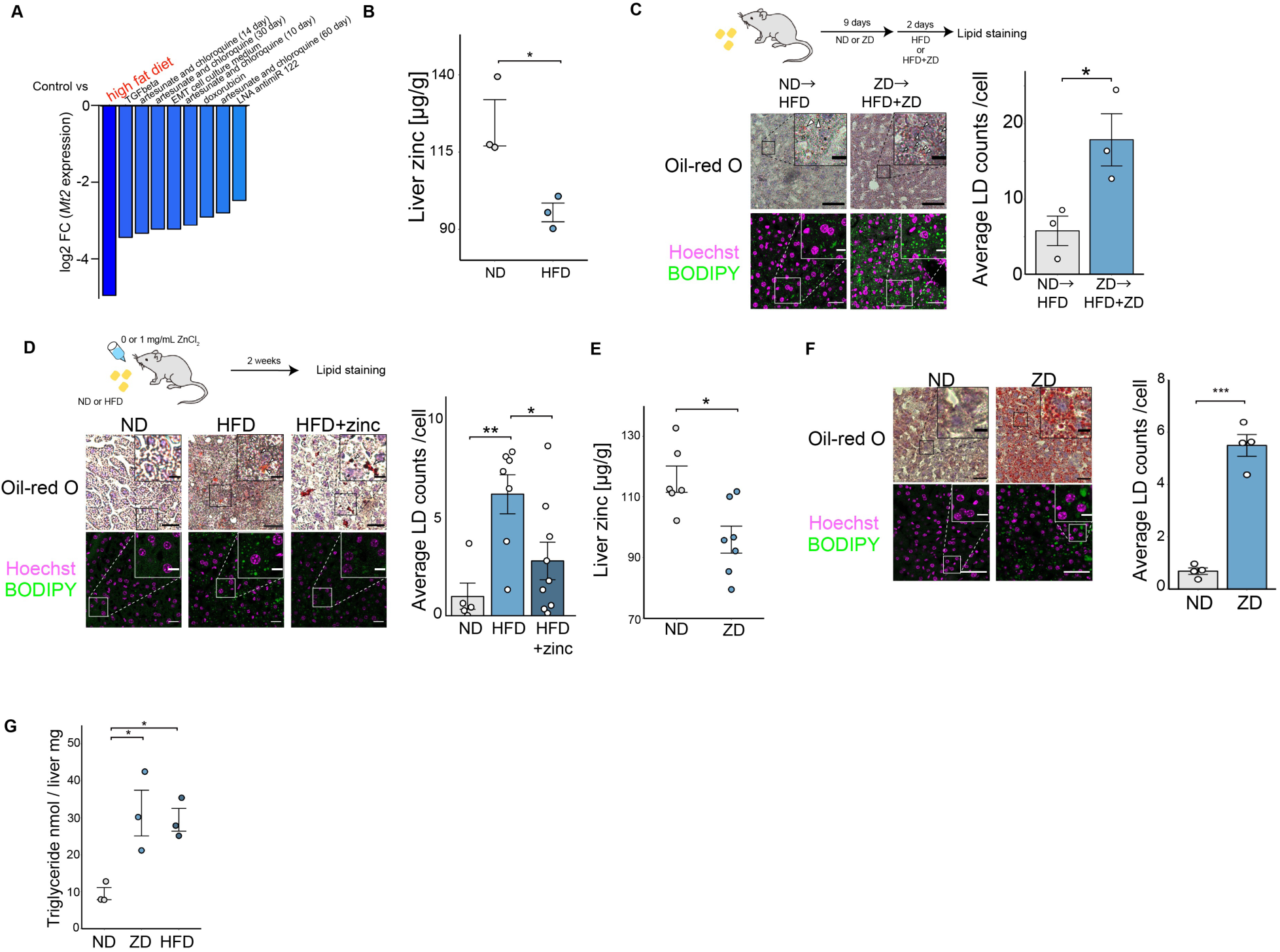
Lipid accumulation in the liver during zinc deficiency A) Log2 (fold change) values of *Mt2* expression under various conditions in mouse tissues. Gene expression data were obtained from the Expression Atlas database. B) ICP‒MS analysis of liver samples from mice fed the indicated diets for two weeks. The data are presented as means ± SEM (n=3). \**P* < 0.05 by Student’s t test. C) Lipid staining analysis of liver samples from mice fed the indicated diet for the indicated periods. The data are representative of three independent experiments. The graph shows means ± SEM (n=3) of the lipid droplets in the BODIPY-stained images. \**P* < 0.05 by Student’s t test. A total of 120–160 cells per image were evaluated. Scale bar, 50 µm (oil red O staining), 30 µm (BODIPY staining), and 10 µm (inset). D) Lipid staining analysis of liver samples from mice fed the indicated diets for two weeks. ZnCl2 (1 mg/mL) was supplemented with drinking water. The data are representative of three independent experiments. The graph shows means ± SEM (ND, n = 5; HFD, n = 7; HFD+zinc, n = 9) of the lipid droplets in the BODIPY-stained images. \**P* < 0.05 and \*\**P* < 0.01 by one-way ANOVA followed by Dunnett’s multiple comparison test. A total of 60–160 cells per image were evaluated. Scale bars, 30 µm (oil red O staining), 30 µm (BODIPY staining), and 10 µm (inset). E) ICP‒MS analysis of liver samples from mice fed the indicated diets for two weeks. The graph presents the mean ± SEM (ND, n=6; ZD, n=7) of the zinc concentration. \**P* < 0.05 by Student’s t test. F) Lipid staining analysis of liver samples from mice fed the indicated diets for two weeks. The graph shows the mean ± SEM (n=4) of the lipid droplets in the BODIPY-stained images. For each image, between 70 and 120 cells were evaluated. \**P* < 0.05 by Student’s t test. G) Triglyceride quantification assay of liver samples from mice fed the indicated diets for two weeks. The graph presents the mean ± SEM (n=3) of the triglyceride concentration. \**P* < 0.05 by one-way ANOVA followed by Dunnett’s multiple comparison test. Scale bar, 50 µm (oil red O staining), 30 µm (BODIPY staining), and 10 µm (inset). ND, normal diet; HFD, high-fat diet; ZD, low-zinc diet; LD: lipid droplet

First, to examine the effect of zinc deficiency on hepatic steatosis, mice fed a normal or zinc-deficient diet were fed a high-fat diet. Comparative analysis revealed a pronounced exacerbation of hepatic lipid accumulation in mice fed the zinc-deficient diet, as demonstrated by oil red O staining and BODIPY staining (Fig. 5C). Conversely, dietary zinc supplementation in mice fed a high-fat diet attenuated hepatic lipid accumulation (Fig. 5D). These findings implicate zinc deficiency as a contributing factor to the development of fatty liver.

To elucidate the mechanisms underlying the contribution of zinc deficiency to hepatic steatosis, we investigated the hepatic characteristics of a zinc-deficient mouse model. Mice fed a zinc-deficient diet presented a reduction in hepatic zinc levels (Fig. 5B, E). This reduction was accompanied by notable hepatic lipid accumulation (Fig. 5F) and elevated triglyceride levels to a similar extent as those observed with a high-fat diet (Fig. 5G). Collectively, these results suggest that diminished hepatic zinc levels facilitate lipid accumulation in the liver, thereby contributing to the development of hepatic steatosis.

### Loss of H3K14ac promotes lipid accumulation in liver tissue via gene expression upregulation

Next, we investigated the mechanism underlying hepatic steatosis induced by zinc deficiency. Mice fed a zinc-deficient diet presented reduced levels of H3K14ac (Fig. 6A, B). Similarly, mice fed a high-fat diet presented reduced H3K14ac levels, whereas zinc supplementation counteracted this reduction (Fig. 6C, S6A). Notably, KAT7 immunoprecipitated from mouse livers fed a zinc-deficient diet presented significantly decreased HAT activity (Fig. 6D). Moreover, the intraperitoneal treatment of mice with the KAT7 inhibitor WM-3835 resulted in lipid accumulation in the liver (Fig. 6E). These results suggest that a reduction in KAT7 activity and the subsequent decrease in H3K14ac levels in the liver may induce hepatic steatosis. To assess whether zinc deficiency in the liver promotes hepatic steatosis in a cell-autonomous manner, we conducted experiments using mouse primary hepatocytes. TPEN treatment of these cells resulted in a decrease in H3K14ac levels and an increase in lipid droplets, effects that were suppressed by TSA treatment (Fig. 6F, G). Furthermore, treatment with WM-3835 or the knockdown of KAT7 led to increased lipid droplet formation (Fig. S6B-E). Taken together, these results suggest that zinc deficiency contributes at least in part to hepatic steatosis through decreased KAT7 activity and a reduction in H3K14ac.

**Fig. 6.**
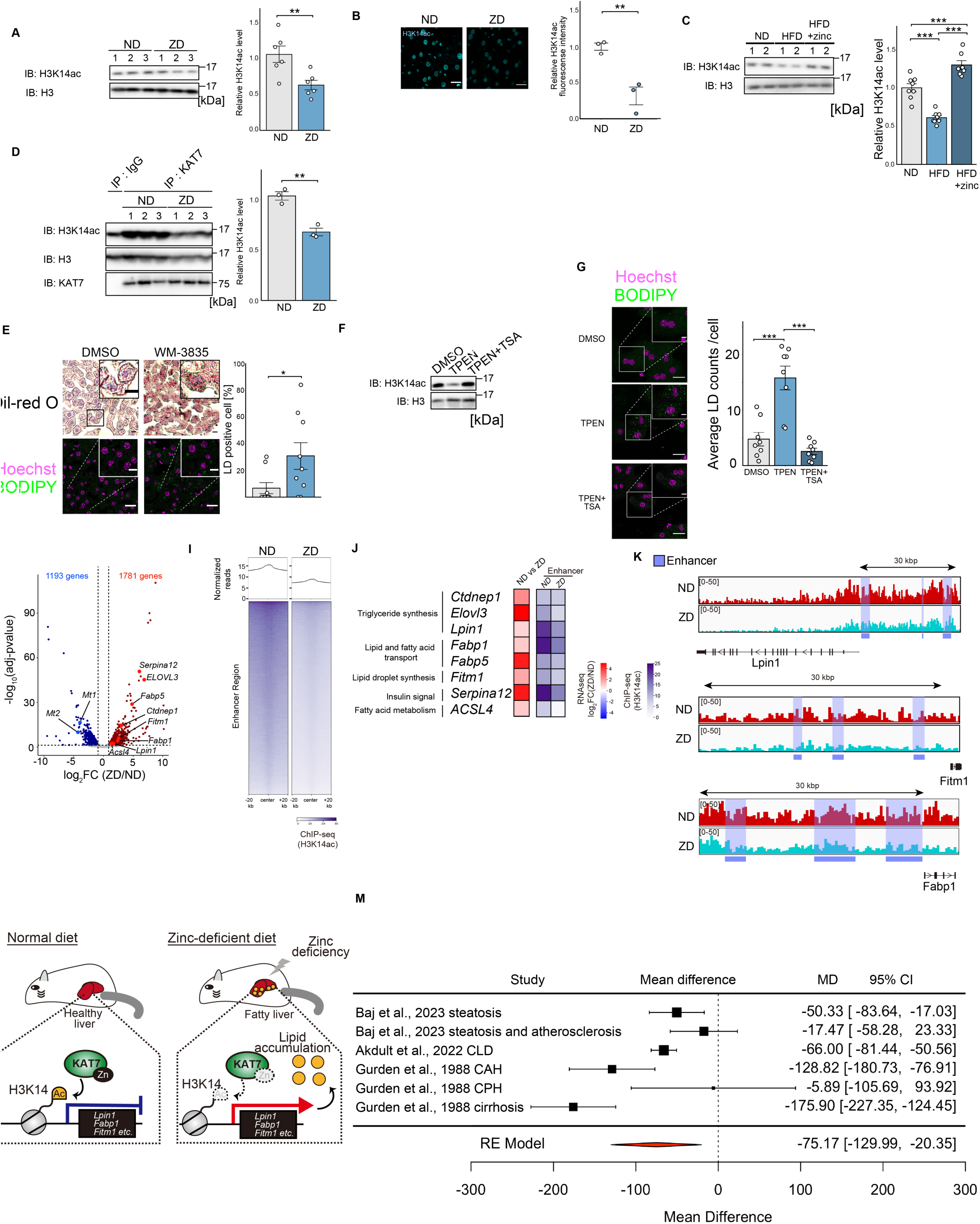
Loss of H3K14ac promotes lipid accumulation in liver tissue via gene upregulation A) Left, Immunoblot analysis of liver samples from mice fed the indicated diets for two weeks. Representative data from six independent mice are shown. Right, Quantification of the H3K14ac immunoblot data. The graph presents the mean ± SEM (n=6) of the average H3K14ac intensity. \*\**P* < 0.01 by Student’s t test. B) Immunofluorescence analysis of liver samples from mice fed the indicated diets for two weeks. The data are representative of three independent experiments. Quantification of H3K14ac fluorescence staining data. A total of 100–150 cells per image were evaluated. The graph presents the mean ± SEM (n=3) of the average H3K14ac intensity. \*\**P* < 0.01 by Student’s t test. C) Immunoblot analysis of liver samples from mice fed the indicated diets for two weeks. ZnCl2 (1 mg/mL) was supplemented with drinking water. Right, Quantification of the H3K14ac immunoblot data. The graph presents the mean ± SEM (ND and ZD, n=8; HFD + zinc, n=7) of the average H3K14ac intensity. \*\*\**P* < 0.001 by one-way ANOVA followed by Dunnett’s multiple comparison test. D) *In vitro* HAT assay. KAT7 immunoprecipitated from liver samples was subjected to an *in vitro* HAT assay. Quantification of the H3K14ac immunoblot data in Fig. 6D. The graph presents the mean ± SEM (n=3) of the average H3K14ac intensity. \*\**P* < 0.01 by Student’s t test. E) Lipid staining analysis of liver samples from mice treated with DMSO or 10 mg/kg WM-3835 for two weeks via daily intraperitoneal injections. The data are representative of three independent experiments. Scale bar, 50 µm (oil red O staining), 20 µm (BODIPY staining), and 10 µm (inset). A total of 15–70 cells per image were evaluated. The graph shows the mean ± SEM (DMSO, n=7; WM-3835, n=9) of the lipid droplets in the BODIPY-stained image. \**P* < 0.05 by Student’s t test. F) Immunoblot analysis of mouse primary hepatocytes treated with the indicated stimuli. The primary hepatocytes were treated with 3 µM TPEN for 10 hours after pretreatment with 0.5 µM TSA for one hour. The data are representative of three independent experiments. G) Lipid staining analysis of mouse primary hepatocytes treated with the indicated stimuli. The primary hepatocytes were treated with 3 µM TPEN for 10 hours after pretreatment with 0.5 µM TSA for one hour. The graph shows the mean ± SEM (n=8) of the lipid droplets. \*\*\**P* < 0.001 by one-way ANOVA followed by Student’s t test followed by Bonferroni post hoc correction. A total of 5–30 cells per image were evaluated. Scale bar, 30 µm (BODIPY staining), 10 µm (inset). H) Volcano plot of the RNA-seq data. Total RNA isolated from liver samples of mice fed the indicated diets for two weeks was used for RNA-seq analysis. I) Average plot (top) and heatmap (bottom) of H3K14ac ChIP-seq reads around enhancer regions (n=76141). J) Heatmaps showing gene expression changes during zinc deficiency and the H3K14ac signal in enhancer regions of the mouse liver. K) Read density of H3K14ac around the genes involved in lipid droplet synthesis in liver samples of mice fed the indicated diets for two weeks. Gene tracks were visualized via the Integrative Genomics Viewer. The data are representative of two independent experiments. L) Schematic of the model reported in this study.M) Meta-analysis of the zinc concentration in the livers of patients with steatosis-related diseases. ND, normal diet; ZD, low-zinc diet; LD: lipid droplet

Next, we investigated whether zinc deficiency in the liver alters the expression of genes associated with lipid accumulation. RNA-seq analysis revealed that zinc deficiency in the liver induced a decrease in the expression of 1,193 genes and an increase in the expression of 1,781 genes (Fig. 6H). We then conducted ChIP-seq analysis for H3K14ac, the results of which revealed a global reduction in H3K14ac at enhancer regions during zinc deficiency (Fig. 6I). Notably, the expression of several genes associated with lipid droplet synthesis pathways, such as triglyceride synthesis, intracellular lipid transport, and lipid droplet budding, was upregulated during zinc deficiency, along with a decrease in H3K14ac at enhancer regions (Fig. 6J, K). Additionally, the treatment of primary hepatocytes with WM-3835 increased the expression of key genes involved in lipid droplet formation (Fig. S6F). Collectively, these findings suggest that zinc deficiency drives lipid accumulation by upregulating the expression of genes involved in lipid droplet synthesis, a process mediated by reduced KAT7 activity and diminished H3K14ac levels (Fig. 6L).

Finally, we conducted a clinical literature review to examine the association between zinc and fatty liver diseases in humans. Several clinical studies have measured hepatic zinc concentrations in both healthy individuals and patients with diseases accompanied by hepatic steatosis. A meta-analysis of these studies revealed significantly lower hepatic zinc levels in the patient group than in the healthy control group (Fig. 6M). This finding suggests that zinc deficiency in the liver may serve as a potential risk factor for fatty liver diseases, including NAFLD.

## Discussion

In this study, we elucidated the molecular mechanism by which zinc deficiency stress is converted to a decrease in H3K14ac. Moreover, we presented examples of genes that are controlled by this decrease in epigenetic signalling, including those related to zinc homeostasis and the accumulation of lipid droplets. We provided evidence that H3K14ac regulates gene expression via its enhancer regions; however, the mechanism remains unclear. H3K14ac acts as a binding site for numerous bromodomain proteins, referred to as “readers”^47^. Our results suggest that transcriptional repressive readers may reside in the enhancer regions of ZIP10 and several genes implicated in lipid droplet synthesis in the basal state. Several groups, including our group, have reported that bromodomain-containing proteins such as ZMYND8 and BAZ2A recognize H3K14ac on enhancer regions to repress gene transcription^36–38^, whereas at least 10 readers have been shown to recognize H3K14ac^48^. The identification of the specific reader that recognizes H3K14ac on the enhancer is a future aim.

The results of our present study suggest that zinc deficiency may lead to a loss of zinc coordination in KAT7; however, how zinc is released from KAT7 remains unclear. Although TPEN is a chelator with a high affinity for zinc^49^, *in vitro* TPEN treatment did not affect the coordination of zinc or the HAT activity of KAT7 (Fig.

S3G, H). This result suggests that zinc is tightly coordinated in KAT7 and that zinc is unlikely to be spontaneously released from KAT7 during zinc deficiency. One possible mechanism is that unidentified cellular molecule(s) may recognize the decreased concentration of zinc and facilitate zinc dissociation from KAT7 in the cell. Another possibility is that the activity of certain chaperones that help coordinate zinc in KAT7 may be reduced under zinc-deficient conditions. Recent reports have demonstrated that proper zinc transport mediated by a zinc chaperone regulates the activity of zinc-dependent enzymes^50,51^. Moreover, *in vitro* zinc repletion of WT KAT7 immunoprecipitated from cells treated with TPEN did not fully restore KAT7 HAT activity (Fig. 3H). This raises the possibility that something irreversible other than the loss of zinc coordination, such as changes in the PTMs of KAT7, occurs in cells under zinc-deficient conditions. Additional studies are necessary to elucidate the precise regulatory mechanism of KAT7 HAT activity during zinc deficiency.

The regulation of ZIPs on the plasma membrane at the protein level contributes to the maintenance of zinc homeostasis. For example, ZIP4 is degraded via endocytosis under normal conditions, whereas endocytosis is suppressed during zinc deficiency, leading to an increase in the amount of ZIP4 on the plasma membrane and an increase in zinc influx^52,53^; another example includes the phosphorylation-dependent regulation of ZIP7 activity^54^. However, little is known about the regulation of ZIP10 at the protein level, except that ZIP10 forms a heteromeric complex with ZIP6^55^. The regulatory mechanisms of ZIP10 at the protein level need to be investigated.

The accumulation of intracellular lipid droplets observed in fatty liver progresses through multiple steps, including triacylglycerol synthesis at the endoplasmic reticulum membrane, lipid budding from the endoplasmic reticulum to the cytoplasm, and the maturation of lipid droplets in the cytoplasm. In this study, we showed that zinc deficiency upregulated the expression of genes involved in these steps, for example, *Lpin1*, *Fabp1*, and *Fitm1*. LPIN1 is an enzyme that catalyses the conversion of phosphatidic acid to diacylglycerol during triglyceride biosynthesis.

Several reports have suggested that the inhibition of LPIN1 ameliorates hepatic steatosis via the suppression of triglyceride biosynthesis^56,57^. FABP1 plays a critical role in the uptake and intracellular transport of fatty acids, and silencing Fabp1 has been shown to reduce lipid accumulation in NAFLD model mice^58^. FITM1 directly binds to triglycerides and induces lipid droplet formation^59^. The overexpression of FITM1 has been shown to increase lipid droplet formation in cultured cells. On the basis of these previous reports, zinc deficiency may promote lipid synthesis at multiple stages by increasing the expression of these genes. However, the specific impact on each stage needs to be investigated further.

The findings of this study suggest that zinc deficiency contributes to hepatic lipid accumulation through a reduction in H3K14ac. On the other hand, lipid accumulation induced by the intraperitoneal injection of the KAT7 inhibitor was relatively mild compared with that caused by zinc deficiency. Previous research has shown that zinc deficiency can trigger oxidative stress and endoplasmic reticulum (ER) stress^16,60^. Since oxidative and ER stress are known to promote lipid droplet synthesis^61,62^, it is plausible that zinc deficiency further exacerbates hepatic lipid accumulation through these mechanisms. Additionally, studies have demonstrated that the knockout of GPR39, an extracellular zinc sensor, induces hepatic lipid accumulation in mice^63^. This phenotype is consistent with the effects observed in zinc deficiency, which may inactivate GPR39. Together, these findings indicate that zinc deficiency may contribute to hepatic lipid accumulation through multiple interconnected pathways.

Although zinc deficiency has been implicated in numerous human diseases, the causal molecular mechanisms remain largely unexplored. This study provides evidence that zinc deficiency is associated with a loss of KAT7 activity, advancing our understanding of the mechanisms underlying various diseases caused by zinc deficiency. For example, zinc deficiency during pregnancy has devastating effects on newborns, such as growth impairment and low birth weight^64^. From the viewpoint of KAT7, *Kat7* knockout mice show decreased transcription of genes essential for embryonic development owing to a global reduction in H3K14ac, which leads to embryonic lethality^65^. These findings suggest a link between zinc deficiency and the loss of KAT7 activity during embryonic development. Moreover, KAT7 plays key roles in immunity by regulating the functions and differentiation of T cells^66,67^, which was observed in model rodents fed a zinc-deficient diet^68^. Thus, the biological phenomena observed during zinc deficiency are better understood from the perspective of the functional regulation of KAT7-catalysed H3K14ac.

## Materials and Methods

A list of the primers and siRNAs used in this study is provided in Table S1.

## Plasmids

The cDNAs encoding human KAT7, BRPF1, BRPF2, BRPF3, JADE1, JADE2, JADE3, ING4, ING5, and MEAF6 were amplified by PCR using KOD One PCR Master Mix -Blue- (TOYOBO, KMM-201) and inserted into pcDNA3/GW with an HA or a FLAG tag (Invitrogen) or into pMAL-c6T (New England Biolabs). The mutated constructs were prepared by PCR-mediated site-directed mutagenesis. A single guide RNA (sgRNA) targeting *KAT7* (5’-AGCCGCCGGCAATGCCGCGA-3’) was designed using CHOPCHOP (https://chopchop.cbu.uib.no). The sgRNA was inserted into pX459 (Addgene, 62988).

## Cell culture

HEK293A cells, U2OS cells, HCT116 cells, and Neuro2A cells were maintained in high-glucose Dulbecco’s modified Eagle’s medium (DMEM) (Sigma‒Aldrich, D5796; Wako, 044-29765). HeLa cells were maintained in low-glucose DMEM (Wako, 041- 29775). Each type of culture medium was supplemented with 10% foetal bovine serum (FBS) (Gibco, 10270-106) and 100 units/ml penicillin G (Meiji Seika, 01028-85). All cell lines were grown in 5% CO2 at 37°C.

## Preparation of zinc-deficient medium

FBS was incubated with Chelex-100 resin (Bio-Rad, 1422832) at a final concentration of 0.05 g/mL at 4°C with rotation. After 12 h, the mixture was centrifuged at 500 × g for 5 min. The supernatant was passed through a Millex-GV 0.22 µM PVDF filter (Millipore, Cat#SLGVR33RS). Cell culture medium supplemented with Chelex-100 resin-treated FBS at a final concentration of 10% was used as zinc-deficient medium.

## Transfection

Plasmid transfection was performed using Polyethylenimine”MAX” (Polysciences, 24765) or Lipofectamine 2000 (Thermo Fisher Scientific, 11668019) according to the manufacturer’s instructions. siRNA-mediated knockdown was performed using Lipofectamine RNAiMAX (Thermo Fisher Scientific, 13778500) at a final siRNA

concentration of 10 or 20 nM by reverse transfection, according to the manufacturer’s instructions.

## Generation of knockout cell lines

To generate *KAT7* knockout HEK293A cells or HeLa cells, cells were transfected with pX459 encoding a sgRNA targeting *KAT7*. Forty-eight hours after transfection using Lipofectamine 2000, the cells were selected by 1.0 µg/mL puromycin (Gibco, A11138- 03) treatment for another 48 h. The selection medium was replaced with fresh standard medium, and the cells were grown for 3 days. Limiting dilution was performed to obtain single-cell clones. The knockout status of the clones was confirmed by immunoblotting.

## Immunoblotting

Cells were lysed in a cell lysis buffer (20 mM Tris-HCl pH 7.5, 150 mM NaCl, 10 mM EDTA, 1% Triton X-100, 5 µg/mL leupeptin, 1 mM phenylmethylsulfonyl fluoride (PMSF) and 2 nM TSA) for 10 min at 4°C. The lysates were sonicated with 2 sets of 10-sec pulses using an ultrasonic homogenizer (SMT, UH-50). The cell debris was cleared by centrifugation at 17,700 × g for 15 min. The supernatant was mixed with a sample buffer (80 mM Tris-HCl pH 8.8, 80 µg/mL bromophenol blue, 28.8% glycerol, and 4% SDS) supplemented with 10 mM DTT and incubated for 3 min at 98°C. The samples were then resolved by SDS–PAGE and transferred onto Immobilon-P PVDF membranes (Millipore, IPVH00010). Blocking was performed with 5% skim milk (Megmilk Snow Brand) in TBS-T (50 mM Tris-HCl pH 8.0, 150 mM NaCl and 0.05% Tween 20) for 30 min at room temperature. After incubation of the membranes with the indicated primary antibodies for 12-36 hours at 4°C and corresponding HRP-linked secondary antibodies for 1-2 hours at room temperature, the chemiluminescent signals enhanced by ECL Select (Cytiva, RPN2235) were detected using a Fusion Solo 7S instrument (M&S Instruments). Quantification of the bands was performed using Fiji software^69^.

## Coimmunoprecipitation analysis

Cell lysates were incubated with anti-DYKDDDDK-tagged antibody beads (Wako, 016-22784) at 4°C for 10 min. The beads were washed with washing buffer 1 (20 mM Tris-HCl pH 7.5, 500 mM NaCl, 5 mM EGTA and 1% Triton X-100) and washing buffer 2 (20 mM Tris-HCl pH 7.5, 150 mM NaCl and 5 mM EGTA). The washed beads were mixed with sample buffer (80 mM Tris-HCl pH 8.8, 80 µg/mL bromophenol blue, 28.8% glycerol, and 4% SDS) supplemented with 20 mM DTT for 5 min at room temperature and then incubated for 3 min at 98°C. The samples were separated by SDS‒PAGE and immunoblotted with antibodies.

## Immunofluorescence analysis

For confocal microscopy, cells were grown on glass coverslips (Matsunami Glass, C015001). For imaging with a high-content imaging system, cells were grown on a 96- well plate (Corning, 353072). The cells were fixed with 4% paraformaldehyde in PBS for 10 min and permeabilized with 0.2% Triton X-100 in PBS for 5 min. After blocking with 1% BSA (Iwai Chemicals, A001) in TBS-T for 1 h, the cells were incubated with the indicated primary antibodies overnight at 4°C. After washing three times with PBS, the cells were incubated with fluorophore-conjugated secondary antibodies at room temperature for 2 h and then with Hoechst 33342 (Dojindo, 346-07951) for 20 min.

After washing three times with PBS, the cells were mounted with Fluoromount (Diagnostic BioSystems, K024) for confocal microscopy. Images were acquired using an LSM880 with Airyscan (Zeiss) with a 63×/1.40 oil immersion objective or a CellInsight NXT HCS system (Thermo Scientific). The contrast and brightness of the images were adjusted using Fiji software. Quantification of the images was performed using Fiji software or CellProfiler^70^.

## Histone purification

HEK293A cells or HEK293A *KAT7* KO cells were incubated in hypotonic lysis buffer for 30 min at 4°C with rotation. After centrifugation at 10,000 × g for 10 min at 4°C, the cell pellets were resuspended in 0.4 N H2SO4 and incubated for 30 min at 4°C with rotation. After centrifugation at 15,100 × g for 10 min at 4°C, the supernatant was transferred to a fresh 1.5 mL tube. Then, trichloroacetic acid (TCA) was added to the supernatant dropwise and incubated for 30 min on ice. After centrifugation at 15,100 × g for 10 min at 4°C, the pelleted histones were washed twice with ice-cold acetone. The pellet histones were air-dried at room temperature and dissolved in TBS at a final concentration of 3 µg/µL.

## *In vitro* HDAC assay

HEK293A cells treated with DMSO or 10 µM TPEN for 1 h or 2 h were lysed in a cell lysis buffer for *in vitro* HDAC assay (20 mM Tris-HCl pH 7.5, 150 mM NaCl, 10 mM EDTA, 1% Triton X-100, 5 µg/mL leupeptin, and 1 mM PMSF) for 10 min at 4°C. Purified histones were mixed with the cell lysates and incubated at 37°C for 0.5, 1 or 2 h with DMSO or 10 nM TSA. Then, the samples were mixed with an equal volume of 2x sample buffer supplemented with 10 mM DTT and incubated for 3 min at 98°C, followed by SDS‒PAGE and immunoblotting.

## *In vitro* HAT assay

To measure the activity of KAT7 immunoprecipitated from cells, HEK293A cells transfected with FLAG-KAT7 WT or mutants were lysed in cell lysis buffer for an *in vitro* HAT assay (20 mM Tris-HCl pH 7.5, 150 mM NaCl, 10 mM EDTA, 1% Triton X-100, 5 µg/mL leupeptin, and 1 mM PMSF). The cell lysates were incubated with anti-FLAG M2-agarose gel (Sigma‒Aldrich, A2220) at 4°C for 5 min. The beads were washed twice with washing buffer 3 (20 mM Tris-HCl pH 7.5, 150 mM NaCl, 1% Triton X-100) and once with TBS. Purified histones at a final concentration of 0.5 µM were mixed with the beads and with 50 µM acetyl-CoA in TBS and incubated for 30 min at 30°C with frequent mixing (1,200 rpm) using a Thermomixer C (Eppendorf).

The samples were mixed with an equal volume of 2x sample buffer supplemented with 10 mM DTT and incubated for 3 min at 98°C, followed by SDS‒PAGE and immunoblotting. To measure the activity of the recombinant KAT7 HAT domain, 5 µM recombinant protein was used. To measure the activity of the KAT7 from mice liver, the organ lysates were incubated with KAT7 antibody at 4 °C for 16 hours. Then, the lysates were incubated with Protein G SepharoseTM 4 Fast Flow (GE healthcare, 17061802) at 4°C for 2 hours.

## Recombinant protein purification

The KAT7 HAT domain (a.a. 336-611) WT or C371A mutants encoded in pMAL-c6T were expressed in *E. coli* BL21 cells. Expression was induced by incubation in LB medium supplemented with 0.3 mM IPTG at 25°C for 1.5 h. The cells were pelleted and resuspended in PBS supplemented with 1% Triton X-100 and 1 mM PMSF. After sonication with 4 sets of 30-sec pulses using an ultrasonic homogenizer, the cell debris was cleared by centrifugation at 26,800 × g for 30 min. The supernatants were incubated with Ni Sepharose 6 Fast Flow (Cytiva, 17531806) at 4°C for 8 h. The beads were then washed twice with PBS and twice with TBS and eluted with a 250 mM imidazole solution in TBS using Poly-Prep Chromatography Columns (Bio-Rad, 7311550). The eluates were loaded onto a Superose 6 Increase 10/300 GL column on an AKTA purifier (GE Healthcare) and dialyzed in TBS overnight at 4°C. The purity and concentration of the final products were estimated by Coomassie blue staining of SDS– PAGE gels.

## qPCR analysis

Total RNA was isolated from cells using Isogen (Nippongene, 319-90211) and reverse transcribed with ReverTra Ace qPCR RT Master Mix with gDNA Remover (TOYOBO, FSQ-301) according to the manufacturer’s instructions. Samples of the qPCR mixture were prepared using the KAPA SYBR Fast qPCR Kit (Kapa Biosystems, KK4602). qPCR was then performed on a LightCycler 96 (Roche) or QuantStudio 1 (Thermo Fisher Scientific) instrument. Data for each mRNA were normalized to that of RPS18.

## ChIP‒qPCR

Chromatin samples for ChIP‒qPCR were prepared using a SimpleChIP Enzymatic Chromatin IP Kit (Cell Signaling Technology, 9003) according to the manufacturer’s instructions. The chromatin samples were incubated with anti-H3K14ac antibody (Abcam, ab52946) or anti-histone H3 antibody (Cell Signaling Technology, 4499) at 4°C for 8 h with rotation. Then, the samples were incubated with Dynabeads Protein G (VERITAS, DB10004) at 4°C for 2 h with rotation. After washing three times with low-salt buffer (1% SDS, 1% Triton-X, 2 mM EDTA, 20 mM Tris-HCl pH 8.0 and 150 mM NaCl) and once with high-salt buffer (1% SDS, 1% Triton-X, 2 mM EDTA, 20 mM Tris-HCl pH 8.0 and 500 mM NaCl), the chromatin was eluted with elution buffer (1% SDS, 10 mM EDTA and 50 mM Tris-HCl pH 8.0) for 30 min at 65°C with frequent mixing (1,200 rpm) using a Thermomixer C. The purified DNA was analysed as described in the “qPCR analysis” section. ChIP-seq data used for designing primers were downloaded from the ENCODE portal^71^ with the following identifier: ENCFF002QJL. Data visualization was performed with Integrative Genomics Viewer (Broad Institute).

## ZnAF2 assay

Recombinant proteins or ZnCl2 solution for normalization was incubated with 10 µM ZnAF-2 (Goryo Chemical, SK2001-01) in TBS for 30 min at room temperature. The fluorescence signal was detected with a Varioskan microplate reader (Ex/Em = 492/515 nm, Thermo Scientific).

## ERSE sequence analysis

ERSE sequences, CCAAT-(N)9-CCAC[A/G], on the human genome GRCh/hg38 were retrieved from the GGGenome ultrafast sequence search browser. Among the sequences obtained (plus strand, 3764 sequences; minus strand, 3874 sequences), 1897 sequences were annotated as sequences on gene promoters by employing HOMER. The number of ERSE sequences on genes that were upregulated during zinc deficiency was counted programmatically. Gene expression profile data analysed by GEO2R were downloaded from the GEO repository (GSE49657, GSE99204, GSE108923, and GSE135873).

Genes with a fold change in expression greater than 2 were considered upregulated.

## EdU assay

HeLa cells were grown on a 96-well plate and analysed with a Click-iT EdU imaging kit (Invitrogen, C10340). The cells were incubated with 10 µM EdU for 1 h, fixed with 4% paraformaldehyde in PBS for 15 min at room temperature, and washed twice with PBS containing 3% BSA. The cells were permeabilized with 0.5% Triton X-100 in PBS for 20 min at room temperature and washed twice with PBS containing 3% BSA. Then, Click-iT reaction cocktail was added and allowed to react for 30 min at room temperature. After washing twice with PBS containing 3% BSA, the cells were incubated with Hoechst 33342 (1:2000) in PBS for 30 min at room temperature. Images were acquired with a CellInsight NXT automated microscope and analysed with CellProfiler.

## Membrane protein biotinylation assay

Cells were incubated with 0.4 mg/mL biotin (EZ-Link Sulfo-NHS-SS-Biotin, Thermo Fisher Scientific, 21331) in PBS(+) (PBS containing 50 mM MgCl2 and 100 mM CaCl2) and washed twice with 150 mM glycine in PBS(+). Cells were lysed as described in the “Immunoblotting” section. The cell lysates were pulled down with streptavidin agarose resin (Thermo Fisher Scientific, 20353). The resin samples were washed twice with washing buffer 1 and once with washing buffer 2. The samples were subjected to immunoblotting.

## Zinc imaging by ZnDA-3H

Zinc measurement with ZnDA-3H was performed as described previously with minor optimization^44,72^. HeLa cells were reverse-transfected with 10 µM siRNA. The cells were then transfected with Halo-NES and cultured for 24 hours. The cells were replated on 35 mmφ glass bottom dishes (Matsunami, D11130H) that were coated with 1% Cellmatrix Type I-C (Nitta Gelatin, KP-4100). After 24 hours, the cells were incubated with zinc-normal medium or zinc-deficient medium for an additional 48 hours. Cells were washed with Chelex-treated HEPES-buffered HBSS (30 mM HEPES-NaOH, pH 7.4, 5.36 mM KCl, 137 mM NaCl), and incubated with serum-free DMEM containing 250 nM ZnDA-3H and 5 nM HTL-TMR for 30 min. After labelling, the cells were incubated for an additional 30 minutes. After incubation, the cells were washed twice with Chelex-treated HEPES-buffered HBSS, and the medium was exchanged for Chelex-treated HBSS buffer. Multichannel time-lapse images were acquired with 4 averages per frame in 30-second intervals. ZnDA-3H or HTL-TMR was excited at 488 nm with an argon laser or at 561 nm with a DPSS laser and detected by a HyD detector (Leica). Images were acquired for 5 minutes, and then the cells were incubated with containing Chelex-treated HBSS 10 µM ZnCl2 for an additional 10 minutes. To obtain additional images under zinc-deprived and zinc-saturated conditions, the cells were incubated with Chelex-treated HBSS containing 10 µM TPEN for 5 minutes, and then, the cells were incubated with Chelex-treated HBSS containing 400 µM ZnCl2 and 2.0 µM zinc pyrithione for an additional 10 minutes. For presentation, the pseudocolour fluorescence ratio images of ZnDA-3H/HTL-TMR were generated using Fiji software. For quantification of the intensity, we used CellProfiler.

## Meta-analysis

A systematic literature search was conducted in multiple electronic databases (PubMed, Web of Science, and Embase) using the predefined search terms “zinc” and “liver disease.” From the identified articles, the means of and variances in the liver zinc content in healthy subjects and patients with liver disease were extracted for analysis.

All the statistical analyses were conducted using R (R Foundation for Statistical Computing, Vienna, Austria). For studies reporting only median and IQR data, the means and variances were estimated using the “estmeansd” package^73^. A multilevel meta-analysis was performed to address possible correlations among several outcome statistics using the “meta” package^74^.

## Animal studies

All the experiments were performed following the experimental protocol approved by the animal ethics committee of the University of Tokyo. We complied with all relevant ethical regulations for animal use. Male and female C57BL/6J mice were used. The mice were maintained in a specific pathogen-free facility, and age-matched mice were used for the experiments. The mice were fed a ND (Zn: 6 mg/100 g, 13% calories from fat; CE-2, Clea Japan), ZD (Zn: 0 mg/100 g, 13% calories from fat; Clea Japan), HFD (Zn: 6 mg/100 g, 60% calories from fat; CE-2, Clea Japan), or HFD+ZD (Zn: 0 mg/100 g, 60% calories from fat; Clea Japan). Mouse livers were perfused and immersed O/N in 4% paraformaldehyde/PBS for fixation and then cleared with 30% sucrose solution. Fixed livers were embedded in CryoMount I (Muto PureChemicals), and 8 μm-thick sections were cut using a cryostat (Leica), followed by oil red O staining and BODIPY staining.

## Oil red O staining

Oil red O stock solution (oil red O (Wako, 154-02072) 0.03 g/10 mL isopropanol) was diluted 3:2 with Milli-Q water to make an Oil red O working solution. Frozen blocks were sectioned using a cryostat (Leica), and the sections were stained with Oil Red O working solution for 20 minutes. The sections were then washed three times for 5 minutes with Milli-Q water and transferred to haematoxylin solution (Sakura, 6187-2) for 1 minute, followed by three washes with Milli-Q water. The sections were mounted with Fluoromount (Diagnostic BioSystems, K024) for confocal microscopy. Images were acquired using a DM2500B (Leica) with a 40× objective.

## BODIPY staining

Mouse primary hepatic cells were seeded on glass coverslips, fixed with 4% paraformaldehyde in PBS for 30 minutes, and incubated with 0.5 μM BODIPY 493/503 (Cayman Chemical, 25892) for 20 minutes. The coverslips were mounted with Fluoromount (Diagnostic BioSystems, K024) for confocal microscopy. Frozen blocks were sectioned using a cryostat (Leica), and the sections were stained with 0.5 μM BODIPY 493/503 for 3 hours. The sections were subsequently washed twice in 1x PBS and then mounted with Fluoromount (Diagnostic BioSystems, K024) for confocal microscopy. Images were acquired using a TCS SP8 confocal microscope with a 63× oil immersion objective. Lipid droplet counts were quantified using CellProfiler software.

## Liver triglyceride quantification

Liver samples ranging from 40–60 mg were washed three times with 1x PBS. The samples were subsequently lysed with 10% NP-40. The livers were then homogenized using a homogenizer. The samples were boiled at 98 °C for 3 minutes with intermittent vortexing, cooled to room temperature, and then boiled again for complete solubilization of triglycerides. The samples were centrifuged at 10,000 × g for 10 minutes at 4 °C, after which the supernatant (including the lipid layer) was collected. The samples were diluted 1:20 in water for triglyceride measurements. Liver triglycerides were quantified using a triglyceride assay kit (Abcam, ab653363) in accordance with the manufacturer’s instructions.

## ICP‒MS

The zinc concentration in liver tissues was measured via ICP‒MS (Agilent 7800, Agilent Technologies). Liver tissues were dried at 100 °C for 16 hours. Thirty milligrams of dried liver was decomposed using concentrated nitric acid at 150 °C for 15 hours. The decomposed product was dissolved in 0.08 M nitric acid. Zinc (*m/z* 66) was used to measure the zinc concentration, and indium (*m/z* 115) was used as an internal standard to correct the measurement results.

## Primary hepatocyte isolation

Primary hepatocytes were isolated from male and female mice as previously described^75^. Briefly, primary hepatocytes were isolated by two-step collagenase perfusion. After isolation, the cells were resuspended in low-glucose DMEM supplemented with 5% FBS and 100 units/mL penicillin G (Meiji Seika) and then seeded in a six-well plate precoated with rat collagen type I (Sigma, C3867). The culture medium was changed to maintenance medium after three hours. The maintenance medium contained William’s E medium (Gibco, 12551-032), 1% L-glutamine (Wako, 076-00521), and 1% penicillin G. After an overnight incubation, the hepatocytes were used for subsequent experiments.

## RNA-seq

Total RNA isolated by ISOGEN was used for library construction for RNA-Seq analysis. Library preparation was conducted using an NEBNext® Single Cell/Low Input RNA Library Prep Kit for Illumina® (NEB, E6420). Deep sequencing was performed on the Illumina NextSeq2000 platform to obtain 36-bp paired-end reads. Approximately 15 million sequences were obtained and mapped to the reference human genome (hg38) or mouse genome (mm10) via HISAT2 (v.2.2.1). Transcript read counts were determined using featureCounts (v.2.0.6). Differential gene expression analysis was performed using the R (v.4.3.2) package DESeq2 (v.1.42.1).

## ChIP-seq

Chromatin samples were prepared using a SimpleChIP Enzymatic Chromatin IP Kit following the manufacturer’s instructions. For normalization, the samples were mixed with spike-in chromatin (Active Motif, 53083). The chromatin samples were then incubated with an anti-H3K14ac antibody (Abcam, ab52946) and a spike-in antibody (Active Motif, 61686) for 8 hours at 4 °C with rotation. The samples were subsequently purified as described in the “ChIP‒qPCR” section. Library preparation was conducted using an NEBNext® Single Cell/Low Input RNA Library Prep Kit for Illumina® (NEB, E6420). Deep sequencing was performed on the Illumina NextSeq2000 platform to obtain 36-bp paired-end reads. Approximately 30 million sequences were obtained and mapped to the reference human genome (hg38) or mouse genome (mm10) via HISAT2 (v.2.2.1). The H3K14ac signals on the enhancers were calculated programmatically. BED files for enhancer regions were downloaded from the EnhancerAtlas2.0^41^ and FANTOM5^76,77^ databases. The H3K14ac signals on the enhancer regions were analysed via wigglescout, an R package (https://github.com/cnluzon/wigglescout/) on a supercomputer. The supercomputing resource was provided by the Human Genome Center (the Univ. of Tokyo) and the Institute of Statistical Mathematics.

## Quantification and statistical analysis

All the statistical data were analysed via R (ver. 4.0.5) with RStudio (ver. 2022.12.0+353). *P* values < 0.05 were considered to indicate statistical significance. Detailed statistical information for each experiment, including the number of biological replicates (*n*), is shown in the figures and corresponding legends.

## Author contributions

Conceptualization, T.F.;

Methodology, T.F, T.K., T.M., S.M., Y.K., M.S., and H.N.;

Investigation, T.F., S.T., and L.M.; Formal analysis, T.F.; Visualization, T.F. and S.T.; Funding acquisition, T.F. and H.I.; Supervision, T.F. and H.I.;

Writing – Original Draft, T.F. and S.T.;

Writing – Review & Editing, S.T., L.M., T.K., T.M., S.M., I.N., and H.I.

## Code availability

The code used in this study is freely available on GitHub. (https://github.com/FujisawaGroup/zinc_h3k14ac).

## Acknowledgements

We thank all members and ex-members of the Laboratory of Cell Signaling for fruitful discussions. We are grateful to Dr. Amagai, Y (Kyushu University) for helpful discussions, particularly on zinc imaging. We are grateful to Mrs. Takeyama, R (the University of Tokyo) for helpful of ICP-MS operation. We are grateful to Dr. Sugishita, H (the University of Tokyo) for assisting us with RNA-seq and ChIP-seq. We thank the One-stop Sharing Facility Center for Future Drug Discoveries at the University of Tokyo for the use of the Illumina NextSeq2000 and the TCS SP8 confocal microscope. This study was supported by the Japan Agency for Medical Research and Development (AMED) for the Project for Elucidating and Controlling Mechanisms of Aging and Longevity (grant number JP21gm5010001, 23gm0010009s0102, and 23gm1710001s0102 to H.I.), by the Japan Society for the Promotion of Science (JSPS) for the Grants-in-Aid for Scientific Research (KAKENHI; grant numbers JP22K06610 to T.F. and JP21H04760 to H.I.) and the Grant-in-Aid for Scientific Research on Innovative Areas (KAKENHI; grant number JP22H04804 to T.F. and 19H05771 to M.S.), by the Japan Science and Technology Agency (JST) for Moonshot R&D– MILLENNIA Program (grant number JPMJMS2022-18 to H.I.), by the ISM Cooperative Research Program (2024-ISMCRP-2013) (to T.F.), by the researcher exchange promotion program of ROIS (Research Organization of Information and Systems) (to T.F.).

---

**Fig. S1.**
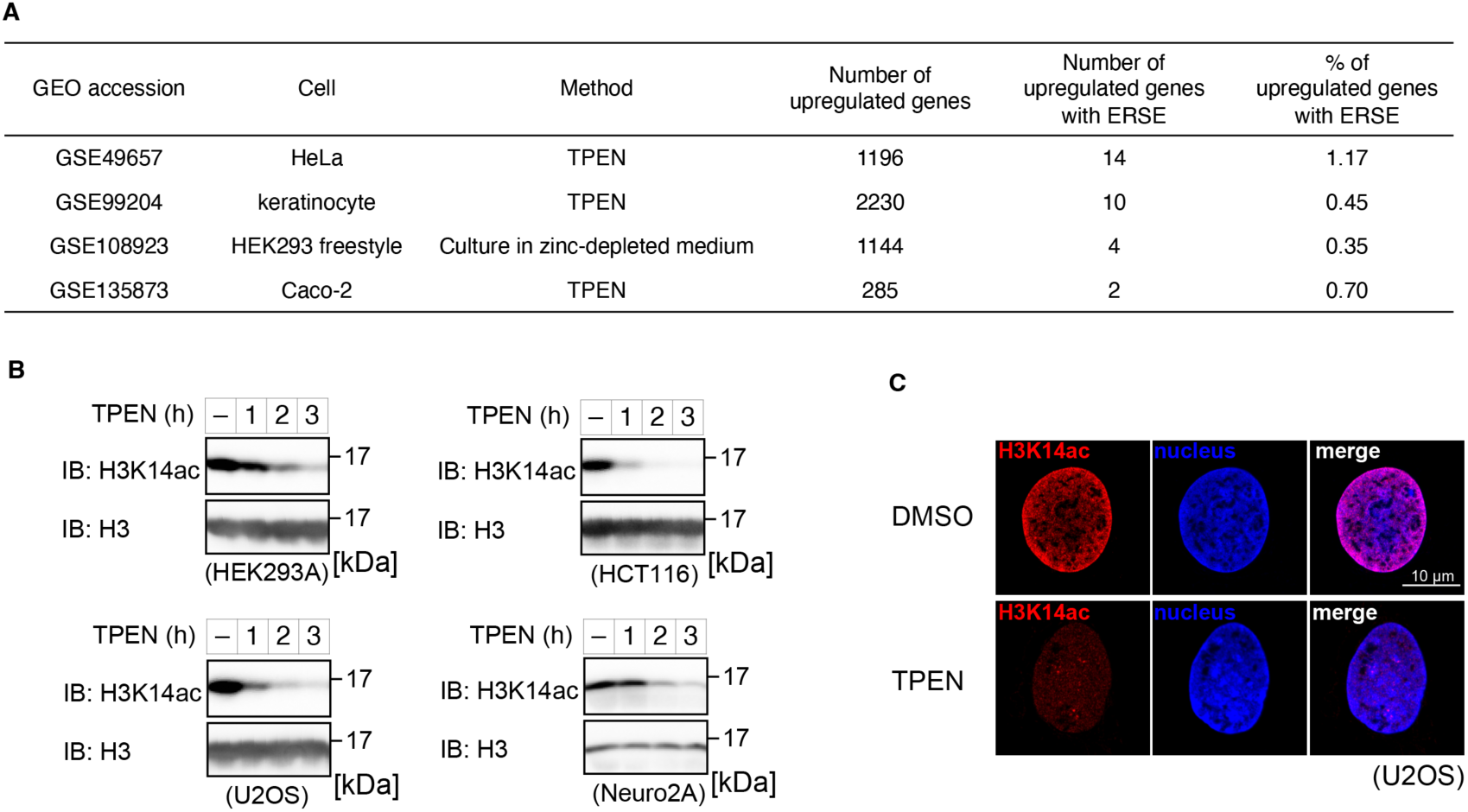
Global decrease in histone H3K14ac during zinc deficiency A) Analysis of the presence of the ATF6-interacting motif (ERSE) in the promoters of the genes whose expression was upregulated under zinc-deficient conditions. B) Immunoblot analysis of cells treated with 10 µM TPEN for the indicated times. The data are representative of three independent experiments. C) Immunofluorescence analysis of U2OS cells treated with 10 µM TPEN for two hours. The data are representative of three independent experiments. Thirty-two (DMSO) or 49 (TPEN) cells pooled from *n =* 3 experiments were analysed. Scale bar, 10 µm.

**Fig. S2.**
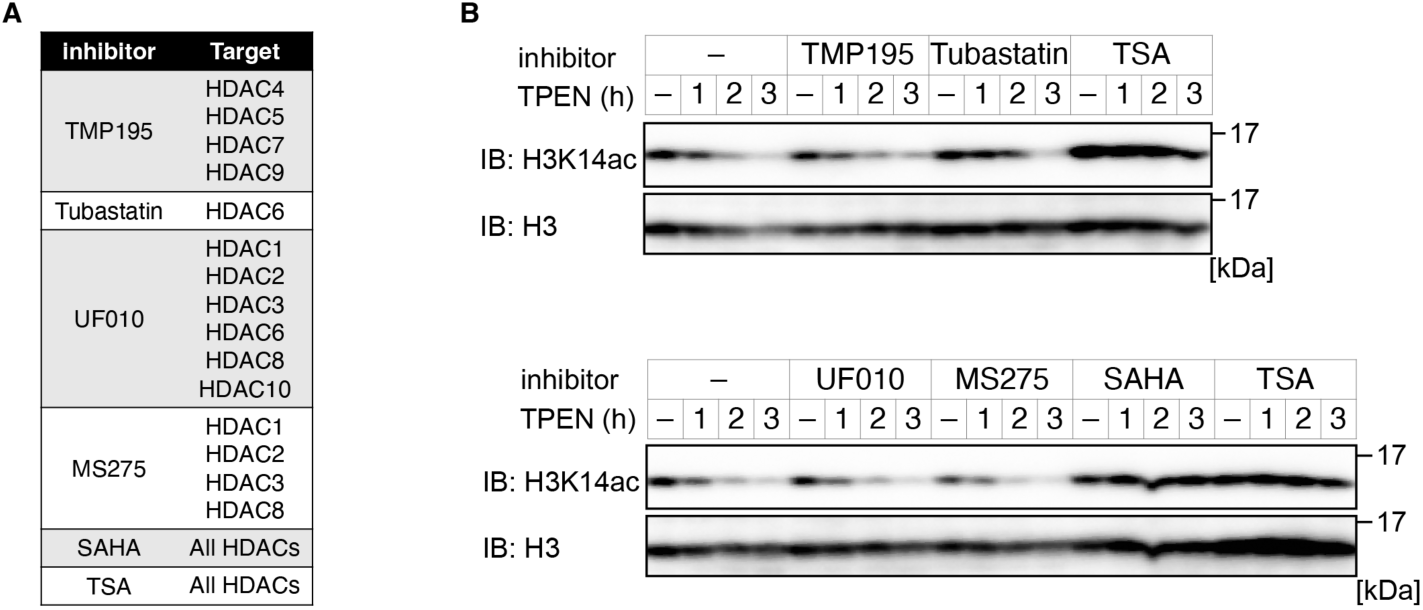
HDACs deacetylate H3K14ac during zinc deficiency A) Classic HDAC family members and their inhibitors. B) Immunoblot analysis of HEK293A cells treated with 10 µM TPEN for the indicated times after pretreatment with HDAC inhibitors for 30 minutes. Representative results of three independent experiments are shown.

**Fig. S3.**
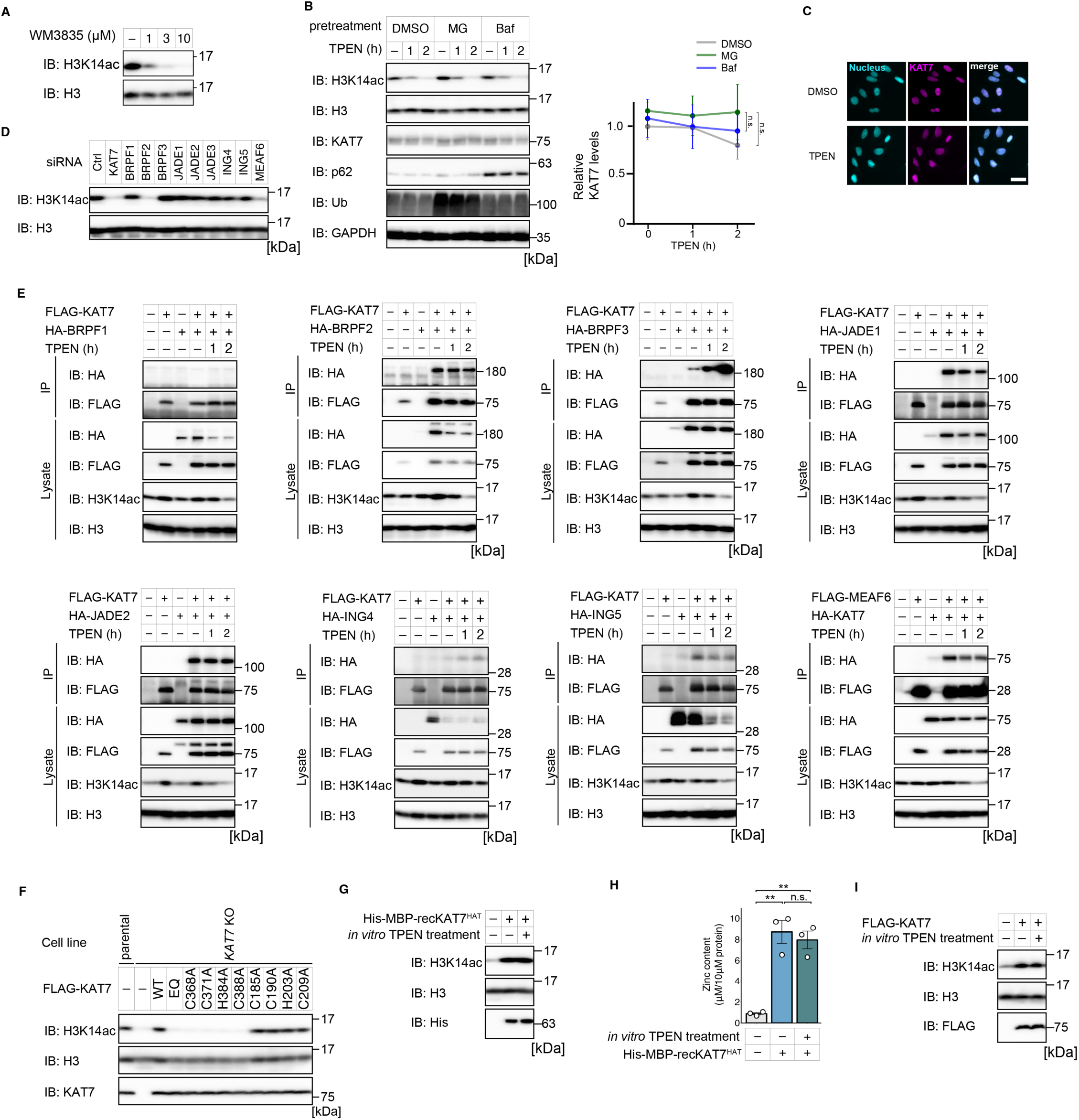
Zinc coordination in the KAT7 MYST domain regulates HAT activity towards H3K14 A) Immunoblot analysis of HEK293A cells treated with WM-3835 at the indicated concentrations for 24 hours. The data are representative of three independent experiments. B) Immunoblot analysis of HEK293A cells treated with 10 µM TPEN for the indicated times after pretreatment with 1 µM MG132 (MG) or 50 nM bafilomycin A1 (Baf) for one hour. Immunoblotting for Ub was performed to determine the effect of MG132. Immunoblotting for p62 was performed to determine the effect of bafilomycin. The data are representative of three independent experiments. The graph shows the mean ± SEM of the KAT7 level normalized to that of GAPDH. The value of the KAT7 level after pretreatment with DMSO was normalized to 1. The results of statistical analyses comparing the reference group (pretreatment with DMSO or two-hour TPEN) to other groups at the same time points are shown. One-way ANOVA with Student’s t test followed by Bonferroni post hoc correction was used. n.s., not significant. C) Immunofluorescence analysis of U2OS cells treated with 10 µM TPEN for two hours. The data are representative of three independent experiments. A total of 142 (DMSO) or 145 (TPEN) cells pooled from *n =* 3 experiments were analysed. Scale bar, 10 µm. D) Immunoblot analysis of HEK293A cells treated with the indicated siRNA for 48 hours. The data are representative of three independent experiments. E) Coimmunoprecipitation analysis showing the interaction between KAT7 and the indicated protein. HEK293A cells were transfected with the indicated plasmids. After 10 µM TPEN treatment for the indicated times, the samples were subjected to coimmunoprecipitation analysis. Data showing the interaction between KAT7 and JADE3 are not included because of the low expression of JADE3. The data are representative of three independent experiments. F) Immunoblot analysis of HEK293A cells and HEK293A *KAT7* KO cells transfected with the indicated plasmids. The data are representative of three independent experiments. G) *In vitro* HAT assay. RecKAT7^HAT^ was incubated with or without 10 µM TPEN for two hours *in vitro*. The samples were subsequently subjected to a HAT assay. The data are representative of three independent experiments. H) ZnAF-2 assay. The zinc content in recKAT7^HAT^ treated with 10 µM TPEN for two hours *in vitro* was measured. The data are presented as means ± SEM (n=3). \*\**P* < 0.01 by one-way ANOVA with Student’s t test followed by Bonferroni post hoc correction. n.s., not significant. I) *In vitro* HAT assay. FLAG-KAT7 WT immunoprecipitated from HEK293A cells was incubated with or without 10 µM TPEN for two hours *in vitro*. The samples were subsequently subjected to the HAT assay. The data are representative of three independent experiments.

**Fig. S4.**
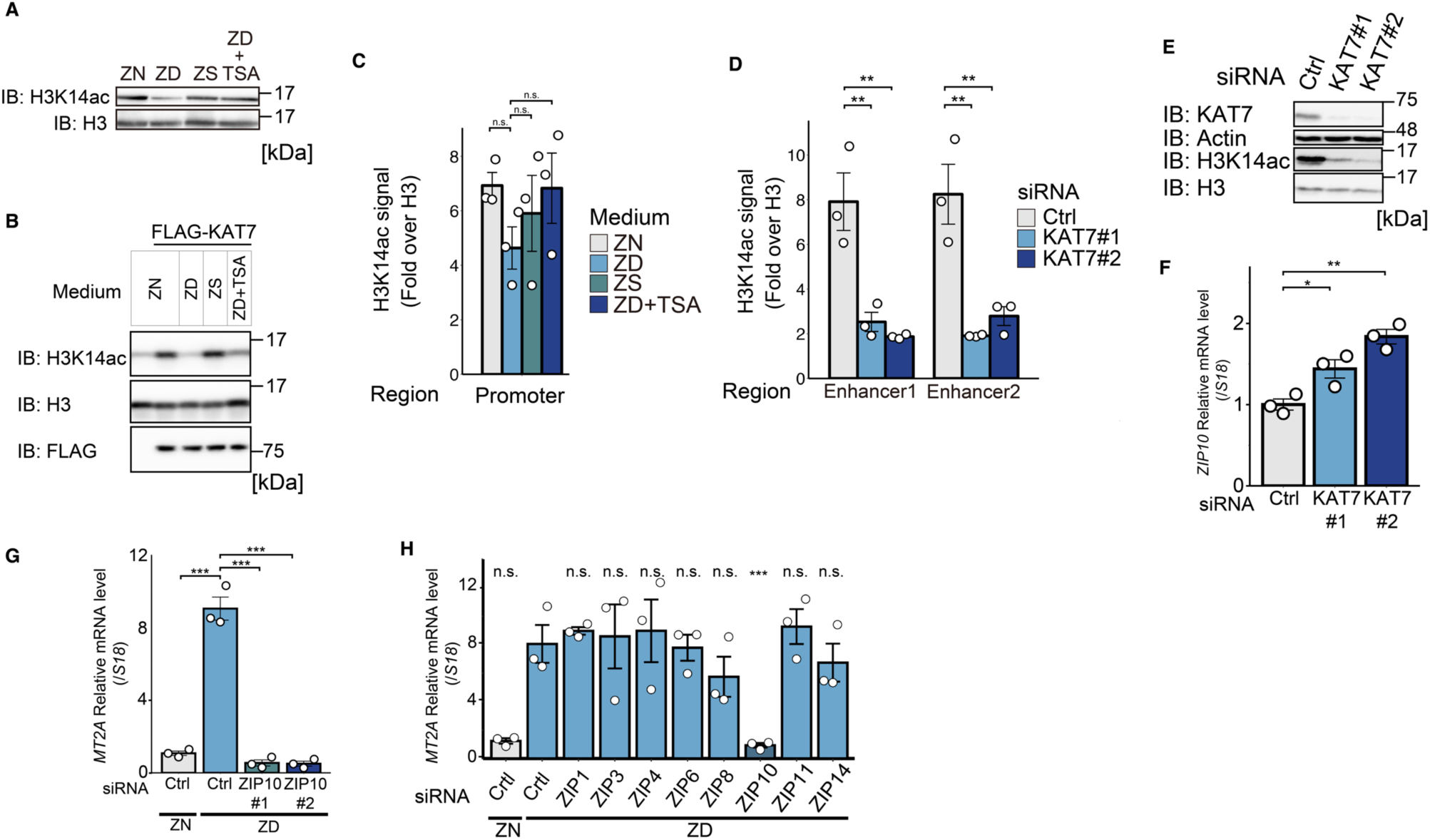
Loss of H3K14ac on the enhancer induces *ZIP10* expression to maintain cellular zinc homeostasis A) Immunoblot analysis of HeLa cells cultured or treated with the indicated conditions. TSA was added every 12 hours. The data are representative of three independent experiments. B) *In vitro* HAT assay. FLAG-KAT7 WT immunoprecipitated from HeLa cells cultured or treated under the indicated conditions was subjected to an *in vitro* HAT assay. TSA was added every 12 hours. The data are representative of three independent experiments. C) ChIP‒qPCR analysis of HeLa cells cultured in the indicated medium for 48 hours. TSA was added every 12 hours. The data are presented as means ± SEM (n=3). n.s., not significant. D) ChIP‒qPCR analysis of HeLa cells transfected with the indicated siRNA for 48 hours. The data are presented as means ± SEM (n=3). \*\**P* < 0.01 by one-way ANOVA with Dunnett’s multiple comparison test. E) Immunoblot analysis of HeLa cells treated with the indicated siRNAs for 48 hours. The data are representative of three independent experiments. F) Quantitative PCR analysis of HeLa cells treated with the indicated siRNAs for 48 hours. The data are presented as means ± SEM (n=3). \**P* < 0.05 and \*\**P* < 0.01 by one-way ANOVA with Dunnett’s multiple comparison test. G) Quantitative PCR analysis. HeLa cells treated with the indicated siRNAs for 24 hours were cultured in the indicated medium for 48 hours. After additional culture with medium supplemented with 1 µM ZnCl2 for 60 minutes, the samples were subjected to quantitative PCR analysis. The data are presented as means ± SEM (n=3). ZN, zinc-normal medium; ZD, zinc-deficient medium. \*\*\**P* < 0.001 by one-way ANOVA followed by Dunnett’s multiple comparison test. H) Quantitative PCR analysis. HeLa cells were treated with the indicated siRNAs for 48 hours. The cells were cultured in zinc-deficient medium for 48 hours, and after additional culture with medium supplemented with 1 µM ZnCl2 for 60 minutes, the cells were subjected to quantitative PCR analysis. The graph shows the mean ± SEM (n=3). \*\*\**P* < 0.001 by one-way ANOVA followed by Dunnett’s multiple comparison test. The results of statistical analyses comparing the reference group (ZD, Ctrl) to other groups are shown. n.s., not significant.

**Fig. S5.**
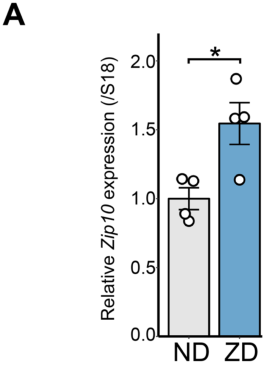
Lipid accumulation in the liver during zinc deficiency A) Quantitative PCR analysis of liver samples from mice fed the indicated diets for one week. The graph shows the mean ± SEM (n=3). \**P* < 0.05 by Student’s t test.

**Fig. S6.**
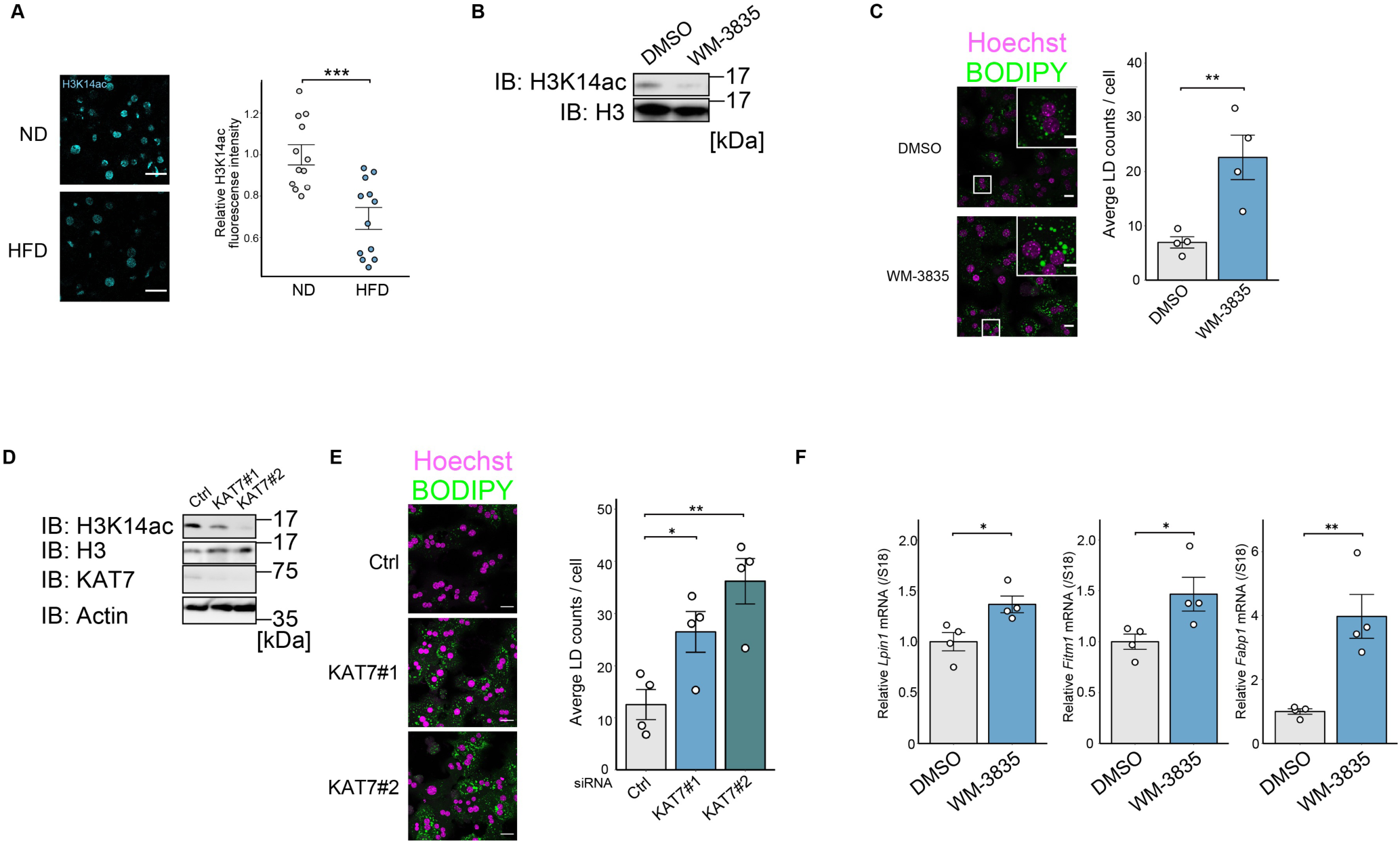
Loss of H3K14ac promotes lipid accumulation in liver tissue via the upregulation of gene expression A) Left, Immunofluorescence analysis of liver samples from mice fed the indicated diets for two weeks. The data are representative of three independent experiments. Scale bar, 20 µm. Right, quantification of H3K14ac fluorescence staining data. For each image, 100–150 cells per image were evaluated. The graph represents the mean ± SEM (n=12) of the average H3K14ac intensity. \*\**P* < 0.01 by Student’s t test. For each image, between 15 and 90 cells were evaluated. B) Immunoblot analysis of mouse primary hepatocytes treated with the indicated stimuli. The data are representative of three independent experiments. C) Lipid staining analysis of mouse primary hepatocytes treated with the indicated stimuli for 24 hours. The graph shows the mean ± SEM (n=3) of the lipid droplets. \**P* < 0.05 by Student’s t test. A total of 20–45 cells per image were evaluated. Scale bar, 50 µm (oil red O staining), 30 µm (BODIPY staining), and 10 µm (inset). D) Immunoblot analysis of mouse primary hepatocytes transfected with the indicated siRNA for 24 hours. The data are representative of three independent experiments. E) Immunofluorescence analysis of mouse primary hepatocytes transfected with the indicated siRNA for 24 hours. A total of 20–45 cells per image were evaluated. The data are representative of three independent experiments. The graph shows the mean ± SEM (n=3) of the lipid droplets. \**P* < 0.05 and \*\**P* < 0.01 by one-way ANOVA followed by Dunnett’s multiple comparison test. F) Quantitative PCR analysis of mouse primary hepatocytes treated with the indicated stimuli. Primary hepatocytes were treated with 1.0 µM WM-3835 for 20 hours. The graph shows the mean ± SEM (n=3). \**P* < 0.05 and \*\**P* < 0.01 by Student’s t test. ND, normal diet; HFD, high-fat diet; LD, lipid droplet

**Table S1.**
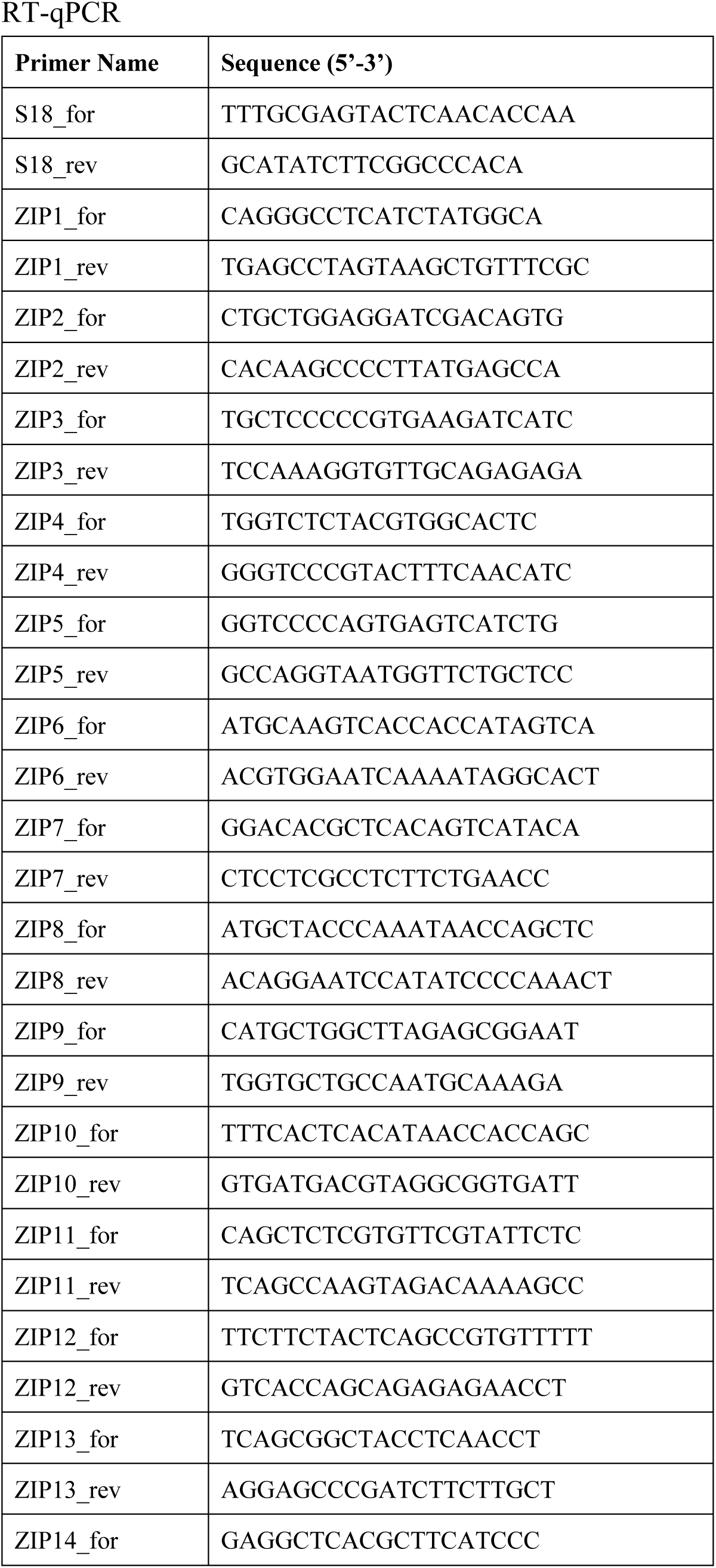
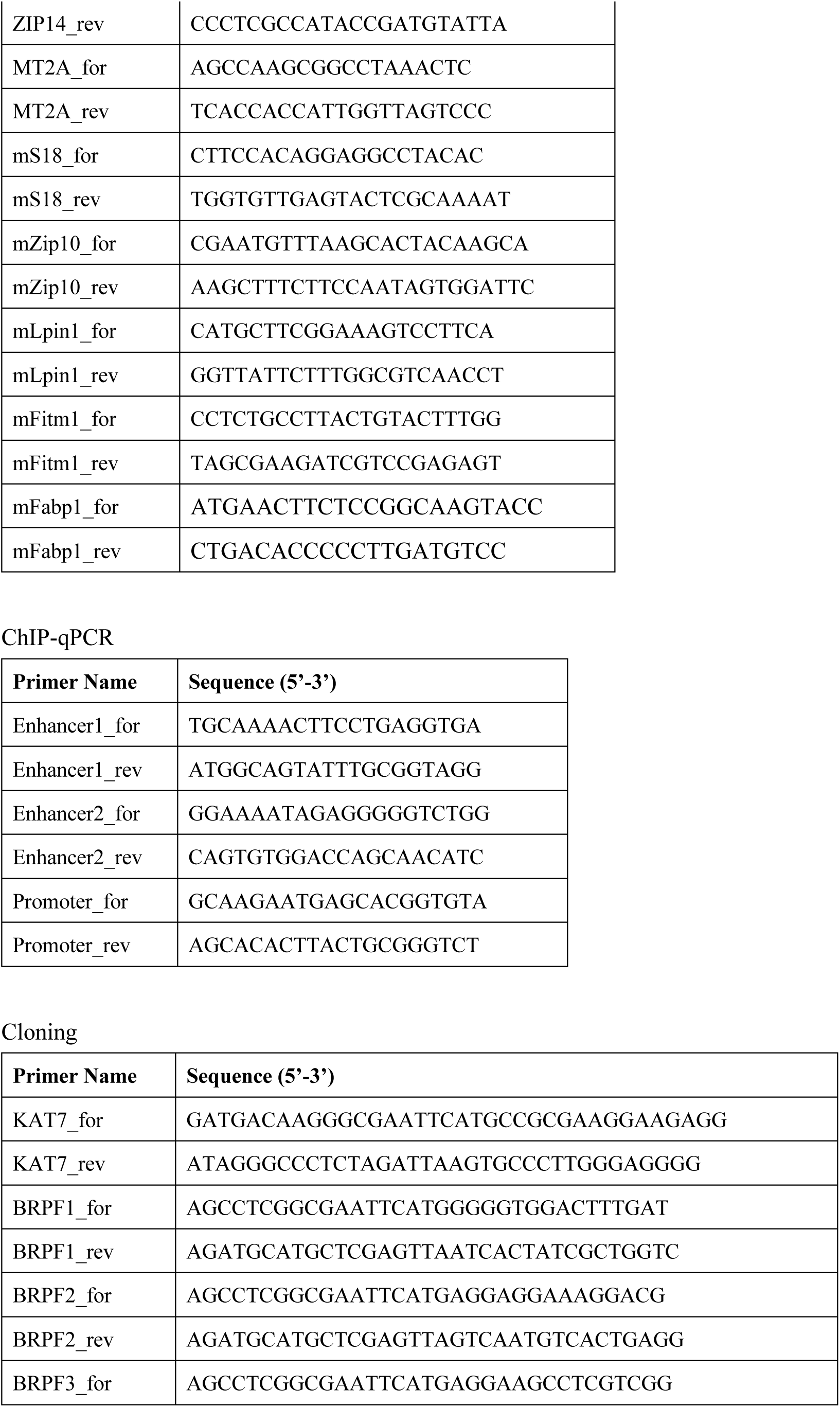
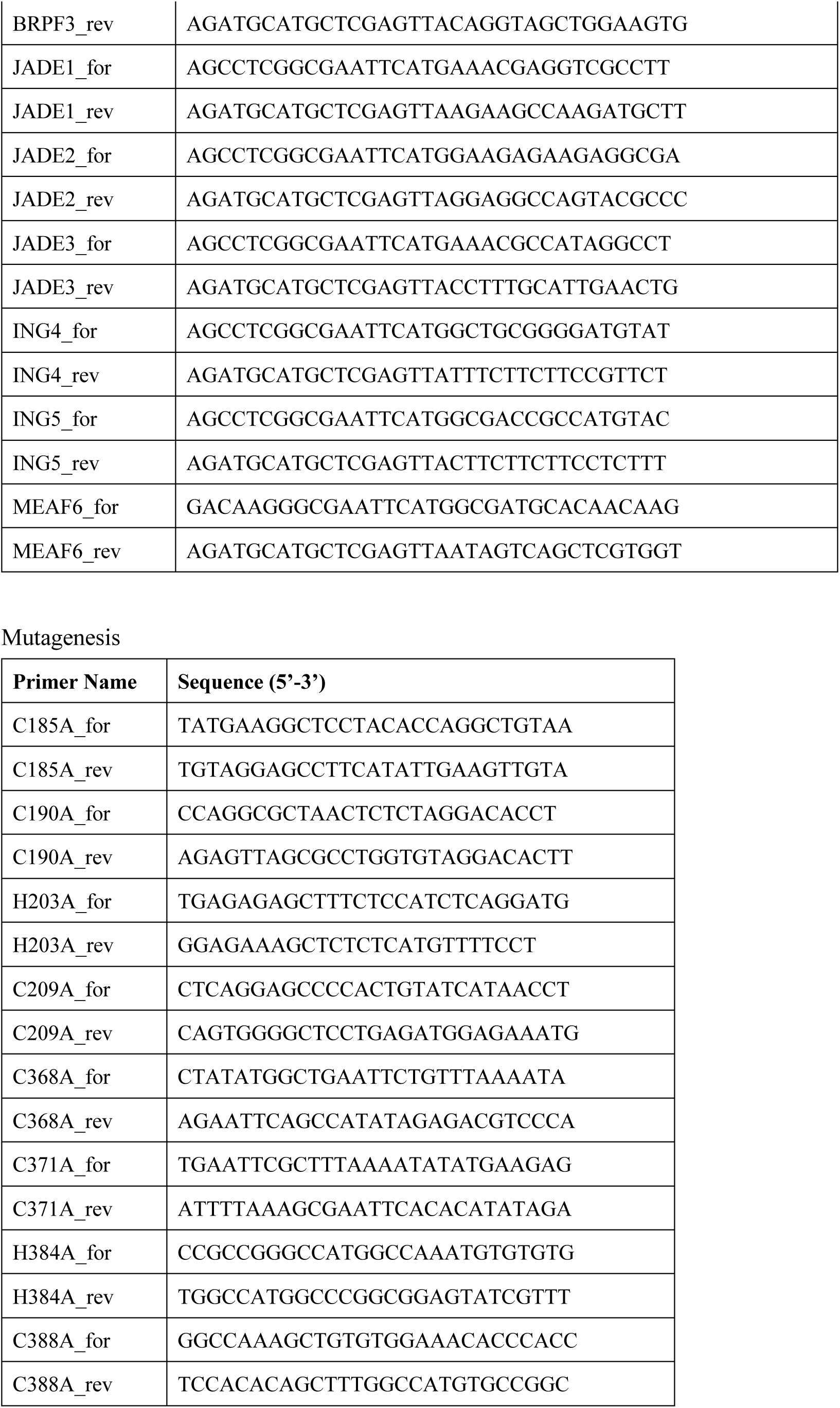
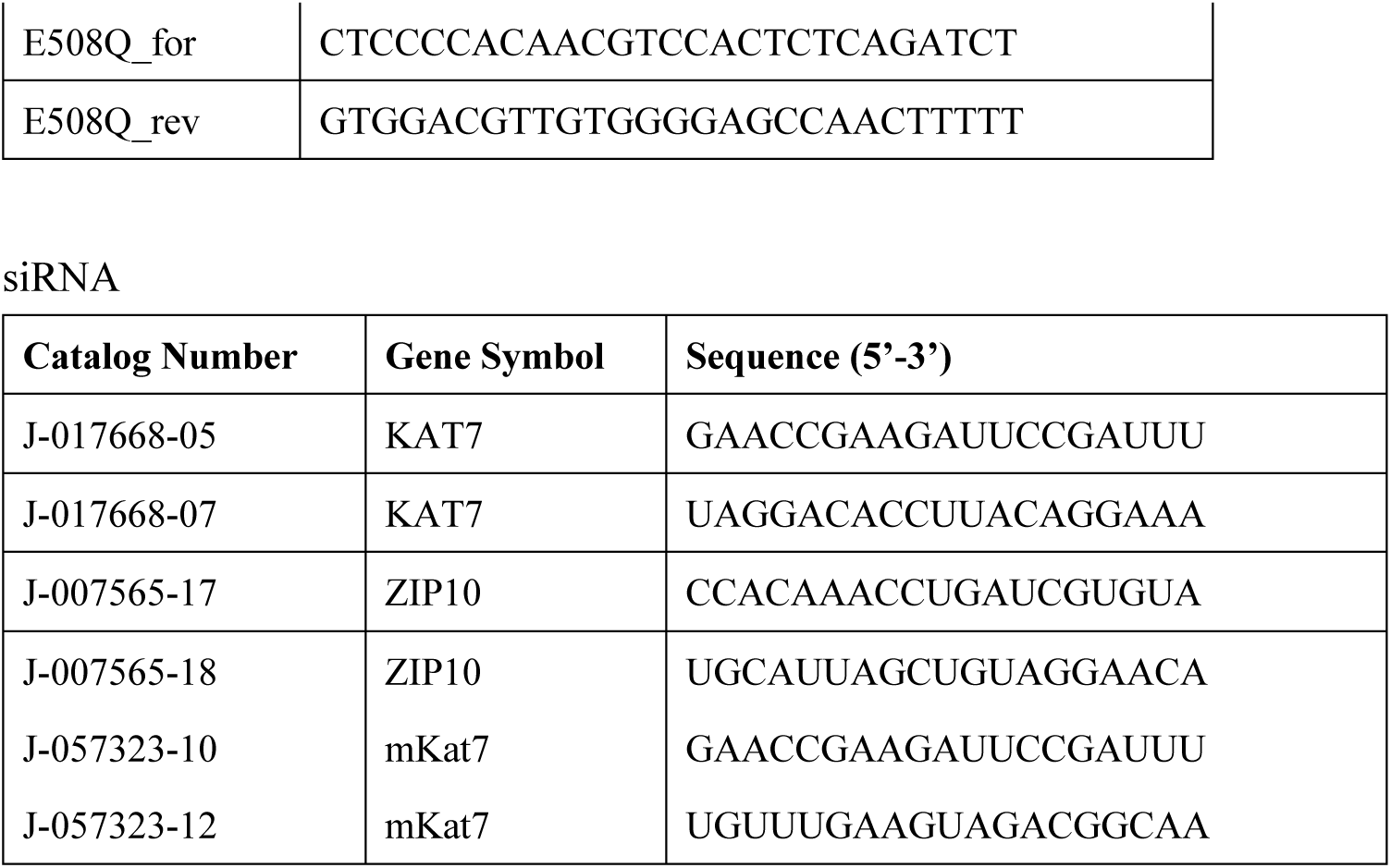
A list of primers and siRNAs used in this study.

